# TDP43 aggregation at ER-exit sites impairs ER-to-Golgi transport

**DOI:** 10.1101/2024.01.24.576891

**Authors:** Hongyi Wu, Loo Chien Wang, Belle M. Sow, Damien Leow, Jin Zhu, Kathryn M. Gallo, Kathleen Wilsbach, Roshni Gupta, Lyle W. Ostrow, Crystal J. J. Yeo, Radoslaw M. Sobota, Rong Li

**Affiliations:** Mechanobiology Institute, National University of Singapore (NUS), Singapore; Functional Proteomics Laboratory, SingMass National Laboratory, Institute of Molecular and Cell Biology, Agency for Science, Technology and Research (A*STAR), Singapore; Department of Anatomy, Yong Loo-Lin School of Medicine, National University of Singapore, Singapore; Department of Neurology, School of Medicine, Johns Hopkins University, USA; Department of Neurology, Lewis Katz School of Medicine, Temple University, USA; National Neuroscience Institute, A*STAR, and Duke-NUS Medical School, Singapore; Lee Kong Chian School of Medicine, Nanyang Technological University, Singapore; Department of Neurology, Feinberg School of Medicine, Northwestern University, USA; School of Medicine, Medical Sciences and Nutrition, University of Aberdeen, UK; Department of Biological Sciences, National University of Singapore, Singapore

## Abstract

Protein aggregation plays key roles in age-related degenerative diseases, but how different proteins coalesce to form inclusions that vary in composition, morphology, molecular dynamics and confer physiological consequences is poorly understood. Here we employed a general reporter based on mutant Hsp104 to identify proteins forming aggregates in human cells under common proteotoxic stress. Over 300 proteins were identified, forming different inclusions containing subsets of aggregating proteins. In particular, TDP43, implicated in Amyotrophic Lateral Sclerosis (ALS), partitions dynamically between two distinct types of aggregates: stress granule and a previously unknown solid inclusion at the ER exit sites (ERES). TDP43-ERES coaggregation is induced by diverse proteotoxic stresses and observed in the motor neurons of ALS patients. Such aggregation causes retention of secretory cargos at ERES and therefore delayed ER-to-Golgi transport, providing a link between TDP43 aggregation and compromised cellular function in ALS patients.

## Introduction

The intracellular space is crowded with proteins of a few hundred gram per liter concentration (Zeskind et al., 2007, Ellis, 2001). To facilitate interaction between partner proteins and prevent unfavorable interaction, a mechanism evolved is the formation of inclusions or aggregates, which assemble functional partners and sequester potentially harmful species such as misfolded proteins (Uversky, 2017, Saad and Jarosz, 2021, Ellis and Minton, 2006, Sontag et al., 2017).

The aggregation of misfolding-prone proteins is a common hallmark of various neurodegenerative diseases, and different proteins aggregate in different diseases (Hipp et al., 2019, Chiti and Dobson, 2006, Soto and Pritzkow, 2018). For example, Aβ and τ are the major constituents of aggregates in Alzheimer’s disease (AD) whereas α-synuclein aggregates are mainly found in Parkinson’s disease (PD). In Amyotrophic Lateral Sclerosis (ALS), various aggregation-prone proteins are implicated, including TAR DNA-binding protein 43 kDa (TDP43), FUS, SOD1, hnRNP A1, UBQLN2 and C9orf72 (Neumann et al., 2006, Soto and Pritzkow, 2018). In addition to their respective hallmark aggregates, secondary aggregate proteins or concomitant proteinopathies are common to neurodegenerative diseases. For instance, Aβ, τ and α-synuclein aggregates all have been implicated in ALS pathology (Moda et al., 2023, Robinson et al., 2018).

Although many proteins can self-aggregate at high concentration when purified *in vitro* (Alberti et al., 2018, Nilsson, 2004), coaggregation of different proteins greatly expands the repertoire of aggregates formed in cells with varied biochemical and physical properties. For instance, a fraction of cytoplasmic TDP43 coaggregates with RNAs, ribosomal proteins and other RNA-binding proteins in stress granules (SGs) and remains soluble (liquid-like) until prolonged stress induces a liquid-to-solid phase transition (Mann et al., 2019, Mateju et al., 2017, Patel et al., 2015). However, when these SGs also contained the nucleoporin NUP62, TDP43 turns insoluble (Gleixner et al., 2022, Chou et al., 2018). Upon proteasome inhibition, many proteins such as CFTR and PS1^A246E^ accumulate ubiquitinated species and aggregate near the nucleus and microtubule organization center (MTOC) to form a single solid inclusion termed aggresome (Johnston et al., 1998, Kopito, 2000). At a similar subcellular location, SOD1^G93A^ coaggregates with either VHL in a liquid-like inclusion termed JUxta Nuclear Quality control compartment (JUNQ), or with polyQ Htt into a distinct solid inclusion termed Insoluble Protein Deposit (IPOD) (Weisberg et al., 2012, Ogrodnik et al., 2014).

The diversity of protein aggregates poses a great challenge for comprehending their physiological consequences, especially given that different aggregates may form at either early or late disease stage and exert cytoprotection or toxicity through distinct mechanisms (Mann et al., 2019, Mateju et al., 2017, Patel et al., 2015). Clinically, aggregates are often categorized by their appearance in patient samples. For example, TDP43 aggregates found in ALS can be classified into skein-like, round, dot-like, linear wisps, and diffuse punctate cytoplasmic staining (DPCS), and it was proposed that DCPS is the earliest type of aggregates found in ALS (Neumann et al., 2006, Kon et al., 2022). However, much remains to be understood about how aggregate morphology relates to the aggregate’s composition, subcellular distribution, biophysical properties, and pathophysiological impacts.

Although many studies have investigated the aggregation of specific disease-associated proteins using protein-specific labels, such as antibodies and fluorescent protein (FP) tags, to observe the formation of diverse protein inclusions requires more general markers that broadly label aggregates of misfolded proteins. In yeast, observation of misfolded protein aggregation often employed Hsp104 tagged with an FP (Erjavec et al., 2007, Liu et al., 2010, Spokoini et al., 2012, Zhou et al., 2011). Hsp104 is a molecular chaperone, a protein disaggregase that recognizes misfolded proteins via interaction with Hsp70/40 chaperones and binding of its N-terminal domain (NTD) to the hydrophobic stretches exposed by misfolded substrates (Grimminger-Marquardt and Lashuel, 2010, Harari et al., 2022). Hsp104 disassembles aggregates by forming hexameric ring-like complexes, through which misfolded proteins are threaded to be disentangled (Yokom et al., 2016, Gates et al., 2017). In yeast, Hsp104-FP foci indicate the locations of aggregates. By using this method, our previous study showed that Hsp104-labelled aggregates are formed on the cytoplasmic surface of ER and mostly remain associated with the ER and mitochondria during their lifetime (Zhou et al., 2014). The human proteome does not appear to contain an Hsp104 ortholog (Shorter, 2011, Saha et al., 2023), but potentiated Hsp104 variants can disaggregate human proteins such as TDP43 and FUS *in vitro* or expressed in yeast (DeSantis et al., 2012, Jackrel et al., 2014), indicating that Hsp104 is capable of recognizing misfolded proteins of human origin.

In this study, we repurposed a mutant Hsp104 that stably binds misfolded proteins as a broad-spectrum reporter/marker for protein inclusions in human cells. Using this reporter, we profiled the protein composition of stress-induced aggregates purified from human cells and uncovered diverse types of aggregates with different but overlapping protein compositions. We observed that TDP43 coaggregates with SEC16A and the COPII protein complex in ER-exit sites (ERES) to form solid inclusions, which morphologically resembles the diffuse punctate cytoplasmic staining (DPCS) of TDP43, the proposed early clinical marker for ALS. Further investigation showed that TDP43-ERES coaggregation impairs ER-to-Golgi transport, suggesting a mechanism for TDP43 to cause cellular dysfunction associated with ALS.

## Results

### Identification of aggregation-prone proteins in human cells using an Hsp104-based reporter

To visualize various types of stress-induced aggregates, we explored the use of FP-tagged Hsp104 as a general marker of protein aggregates in human cells. We introduced the Double Walker B (DWB) mutations (E285Q and E687Q), which fall in the Walker B motifs of both nucleotide-binding domains (NBD1 and NBD2) of Hsp104, to disrupt ATP hydrolysis by Hsp104 required for disaggregation. Hsp104^DWB^ binds but does not disassemble aggregates (Bösl et al., 2005, DeSantis et al., 2012, Jackrel et al., 2014) (Figure S1A and B). To validate its effectiveness as an aggregate marker in human cells, we co-transfected Hsp104^DWB^-mEGFP with aggregation-prone proteins TDP43^Q331K^-mCherry or SOD1^G93A^-mCherry into human RPE-1, a non-transformed cell line that maintains a near-diploid karyotype (Bodnar et al., 1998). Heat stress (42°C), which occurs under certain pathological conditions and destabilizes protein folding (Vabulas et al., 2010, Rzechorzek et al., 2022), was used to exacerbate protein aggregation. Like previous studies, we stressed the cells for a duration of 12 - 16 hr to simulate the late onset of neurodegenerative diseases and allow misfolded proteins to accumulate (Johnston et al., 1998, Kawaguchi et al., 2003, Gamerdinger et al., 2011, Biel et al., 2020). 3D confocal imaging revealed that Hsp104^DWB^ was enriched in most aggregates of TDP43^Q331K^ or SOD1^G93A^, demonstrating that Hsp104^DWB^ recognized aggregates of different proteins (Figure 1A and B). However, over 50% Hsp104^DWB^ aggregates did not colocalize with TDP43^Q331K^ or SOD1^G93A^ aggregates and likely represent aggregates of endogenous proteins (Figure 1A and C).

**Figure 1.**
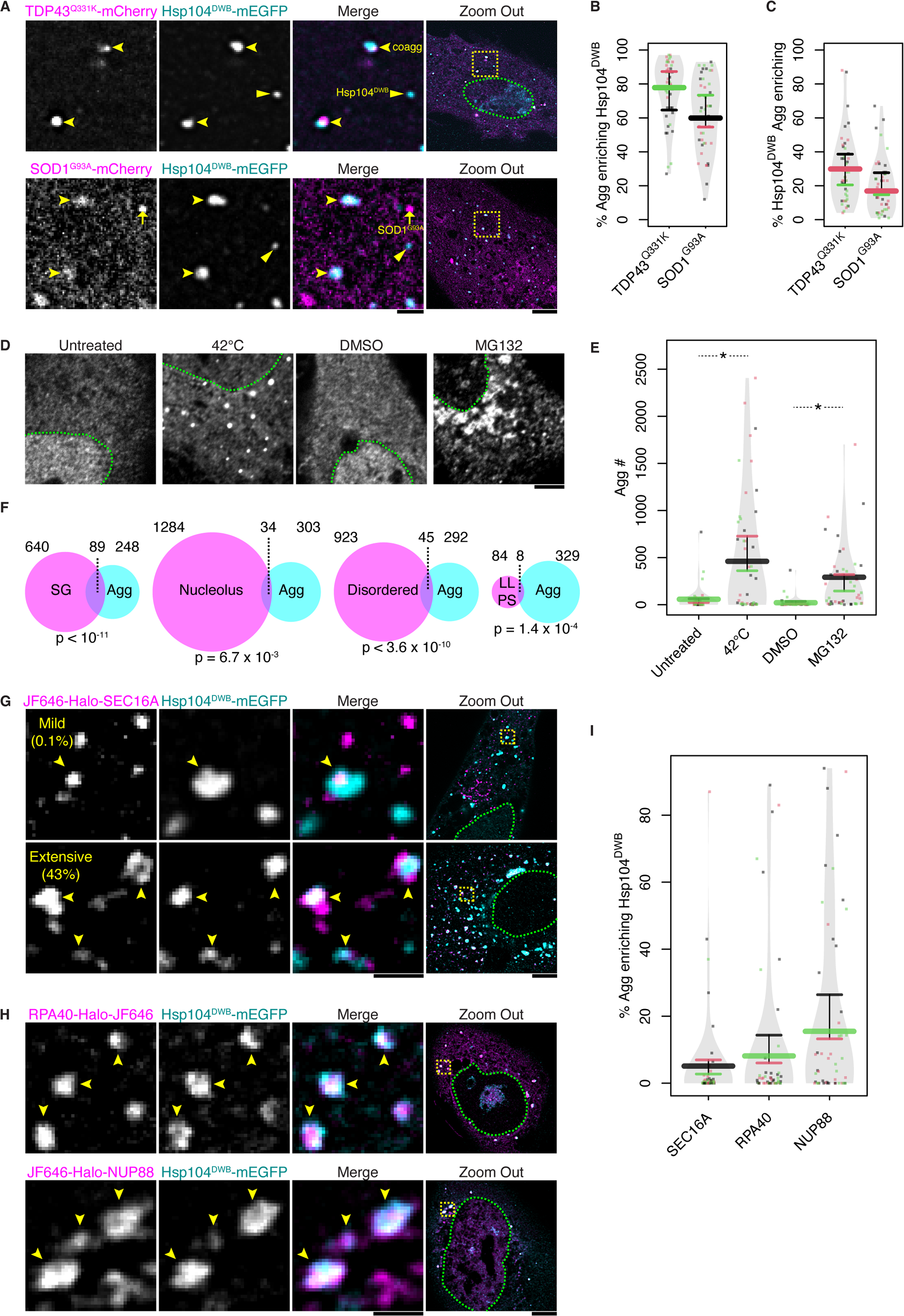
Hsp104^DWB^ enables visualization of aggregates and identification of aggregate proteins in human cells. (**A**) Representative images showing the colocalization between Hsp104^DWB^ and aggregates formed by TDP43^Q331K^ or SOD1^G93A^ after RPE-1 cells were incubated at 42°C for 12 - 16 hr. Representative slices of confocal z-stacks are displayed. Arrowheads indicate Hsp104^DWB^-labelled TDP43^Q331K^ or SOD1^G93A^ aggregates (*i.e.* coaggregate or coagg); arrows indicate SOD1^G93A^ aggregates without Hsp104^DWB^; triangles indicate Hsp104^DWB^ aggregates without TDP43^Q331K^ or SOD1^G93A^. The region displayed in individual channels and Merge views is indicated by the yellow box in “Zoom Out”, and green dashes demarcate the nucleus. Scale bars except in “Zoom Out” represent 1 μm, and represent 5 μm in “Zoom Out”. (**B**) Quantification of the experiments as in (A) showing the percentage (% by number) of TDP43^Q331K^ or SOD1^G93A^ aggregates (Aggs) marked by Hsp104^DWB^. 3 experiments were performed and at least 15 cells were imaged and quantified in each experiment (noted hereafter as N ≥ 3 x 15). The values of individual cells are plotted as dots, and the mean values of cells in the same experiment as horizontal segments, the median of which is elongated and thickened. Different colors represent different experiments. (**C**) Quantification of the experiments as in (A) showing the percentage of Hsp104^DWB^ aggregates that enriched for TDP43^Q331K^ or SOD1^G93A^. N ≥ 3 x 15. (**D**) Representative images showing stress-induced aggregate of endogenous proteins labelled by Hsp104^DWB^-mEGFP in cells without or with heat stress (42°C), or treated with DMSO or 10 μM MG132 for 12 - 16 hr. (**E**) Quantification of the experiments as in (D) comparing the numbers (#) of aggregates in each cell under different conditions. N ≥ 3 x 15. Asterisks (*) indicate p-value ≤ 0.05 in t-tests. (**F**) Venn diagrams showing the overlaps between proteins identified in Hsp104^DWB^-bound aggregates and the protein components of stress granule (SG) and nucleolus and proteins that contain significant disordered regions or undergo *in vitro* liquid-liquid phase separation (LLPS). P-values of Fischer’s exact tests are displayed. (**G**) Representative images of aggregates containing SEC16A and their colocalization with Hsp104^DWB^. Examples for SEC16A inclusions enriching Hsp104^DWB^ mildly (< 10%) and more extensively (≥ 10%) are shown. Arrowheads indicate SEC16A aggregates recognized by Hsp104^DWB^. Experiments hereafter were performed in RPE-1 cells stressed at 42°C for 12 - 16 hr unless indicated otherwise. (**H**) Representative images of the aggregates formed by RPA40 and NUP88 and their colocalization with Hsp104^DWB^. (**I**) Quantification of the experiments as in (G and H) showing the percentage of SEC16A, RPA40 and NUP88 aggregates labelled by Hsp104^DWB^ in each cell. N ≥ 3 x 15.

We subsequently utilized Hsp104^DWB^-mEGFP to observe aggregates of endogenous proteins induced by proteotoxic stress. In untreated or DMSO mock-treated RPE-1 cells, the average number of Hsp104^DWB^ aggregates per cell was close to 0. In contrast, many aggregates labelled with Hsp104^DWB^-mEGFP emerged in cells under heat stress or treatment with the proteasome inhibitor MG132 (Figure 1D and E), another condition often used to induce protein aggregation (Kawaguchi et al., 2003, Gamerdinger et al., 2011). Aggregates enriched for Hsp104^DWB^-mEGFP were also observed in another human cell line, HEK293T, under heat stress (Figure S1C i and ii).

To identify endogenous proteins harbored by stress-induced aggregates, we used Hsp104^DWB^-mEGFP-FLAG as a bait for affinity-purification. This reporter was transfected into human HEK293T cell line, which expresses most proteins found in different tissues and has been widely used for protein purification (Graham et al., 1977, Geiger et al., 2012, Copin et al., 2021). Lysates of heat stressed HEK293T cells were first fractionated by sucrose gradient centrifugation to separate aggregates from soluble proteins (Figure S1C iii - v). Immunoprecipitation was then performed with anti-FLAG tag resin (Figure S1C vi and vii) to further separate aggregates from other macromolecular assemblies and organelles. As a control, cells transfected with Hsp104^DWB^-mEGFP without FLAG tag were subjected to the same workflow. The precipitated proteins were analyzed by liquid chromatography (LC) and quantitative mass-spectrometry (MS) (Figure S1C viii).

337 proteins were found to be enriched in the immunoprecipitate by more than 8.5-fold when compared with the control (Figure S1D, E and Table S1). Gene Ontology (GO) analysis revealed that chaperones and other proteins involved in folding and stress response (*e.g.* Hsp70/40/90 families, chaperonin and VCP) were the top-enriched categories, as expected (Figure S1F, G and Table S1). Moreover, proteins involved in RNA metabolism and translation (*e.g.* TDP43, RNA helicases and translational initiation factors) were overrepresented (Figure S1F, G and Table S1), probably because the prion-like domains present in many RNA-binding proteins promote misfolding and aggregation (Lin et al., 2015, Harrison and Shorter, 2017). Among the top hits there were also nucleoporins, including NUP214, which contains phenylalanine-glycine (FG) repeats, and its close interaction partner NUP88 (Figure S1F, G and Table S1) (Strambio-De-Castillia et al., 2010, Bastos et al., 1997, Bernad et al., 2006). Lastly, in support of the recent findings that macromolecular condensates accumulate misfolded proteins (Mateju et al., 2017, Patel et al., 2015, Frottin et al., 2019), many aggregate proteins found are components of SGs or nucleoli (*e.g.* G3BP1 and RPA40), or contain significant disordered regions, or have been reported to undergo liquid-liquid phase separation (LLPS) *in vitro* (Figure 1F, S1G and Table S1).

Apart from the well-known aggregation-prone proteins such as TDP43 and the SG component G3BP1, we identified many proteins that had not been previously reported to participate in misfolded-protein aggregates. Three candidate proteins that represent different cellular function/components, namely SEC16A, RPA40 and NUP88, were selected for validation by imaging. SEC16A is a cytoplasmic protein that defines ER-exit sites (ERES) by forming liquid-like inclusions at certain locations of the ER surface, and recruits COPII complexes to drive vesicle budding to transport cargos towards the Golgi (Watson et al., 2006, Budnik and Stephens, 2009, Weigel et al., 2021, Gallo et al., 2023). RPA40, also known as POLR1C, is a DNA-dependent RNA polymerase found in the nucleoplasm and the fibrillar centers of nucleoli (Frottin et al., 2019), and NUP88 is a subunit of the cytoplasmic filaments of the nuclear pore complex (Beck and Hurt, 2017). The abundance of these three proteins in aggregates was medium among the identified proteins (Figure S1D and E). Each of these proteins was fused with HaloTag (Halo) and co-transfected with Hsp104^DWB^-mEGFP into cultured cells for confocal imaging.

We first validated in HEK293T cell line that SEC16A, RPA40 and NUP88 were present in Hsp104-labelled aggregates after heat stress (Figure S2A and B). For better observation of subcellular localization, we expressed these proteins in RPE-1 cells, which are bigger in size. Under heat stress, all three proteins formed aggregates, some though not all of which were marked by Hsp104^DWB^-mEGFP (Figure 1G - I). Unlike the aggresome, JUNQ or IPOD, which are aggregates found in the vicinity of the nucleus and MTOC (Johnston et al., 1998, Weisberg et al., 2012, Ogrodnik et al., 2014), Hsp104^DWB^-labelled aggregates of SEC16A, RPA40 or NUP88 were found throughout the cytoplasm (Figure 1G - I). Notably, cells displayed marked heterogeneity in the percentage of aggregates labelled by Hsp104^DWB^. For example, the majority of cells contained no or < 10% SEC16A inclusions recognized by Hsp104^DWB^, whereas a small fraction of cells displayed much more extensive aggregation (> 10% SEC16A inclusions with Hsp104^DWB^) (Figure 1G and I).

### Differential preference in protein coaggregation

We next investigated if the proteins validated above formed the same or different aggregates by analyzing the pairwise coaggregation of these proteins under heat stress. It turned out that not all proteins aggregated into the same inclusions, but each coalesced with its selective partners at different levels of efficiency (Figure 2A – H and S3A). For example, TDP43 coaggregated with G3BP1, SEC16A and NUP88 with sequentially decreasing levels but did not aggregate with RPA40 (Figure 2A - D and H). Again, we observed heterogeneity in the degree of coaggregates (coaggs) formation among cells. For example, the majority of cells co-expressing TDP43-mNG and Halo-SEC16A contained no or low numbers of TDP43/SEC16A coaggs, whereas some cells exhibited much more extensive coaggregation (Figure 2B).

**Figure 2.**
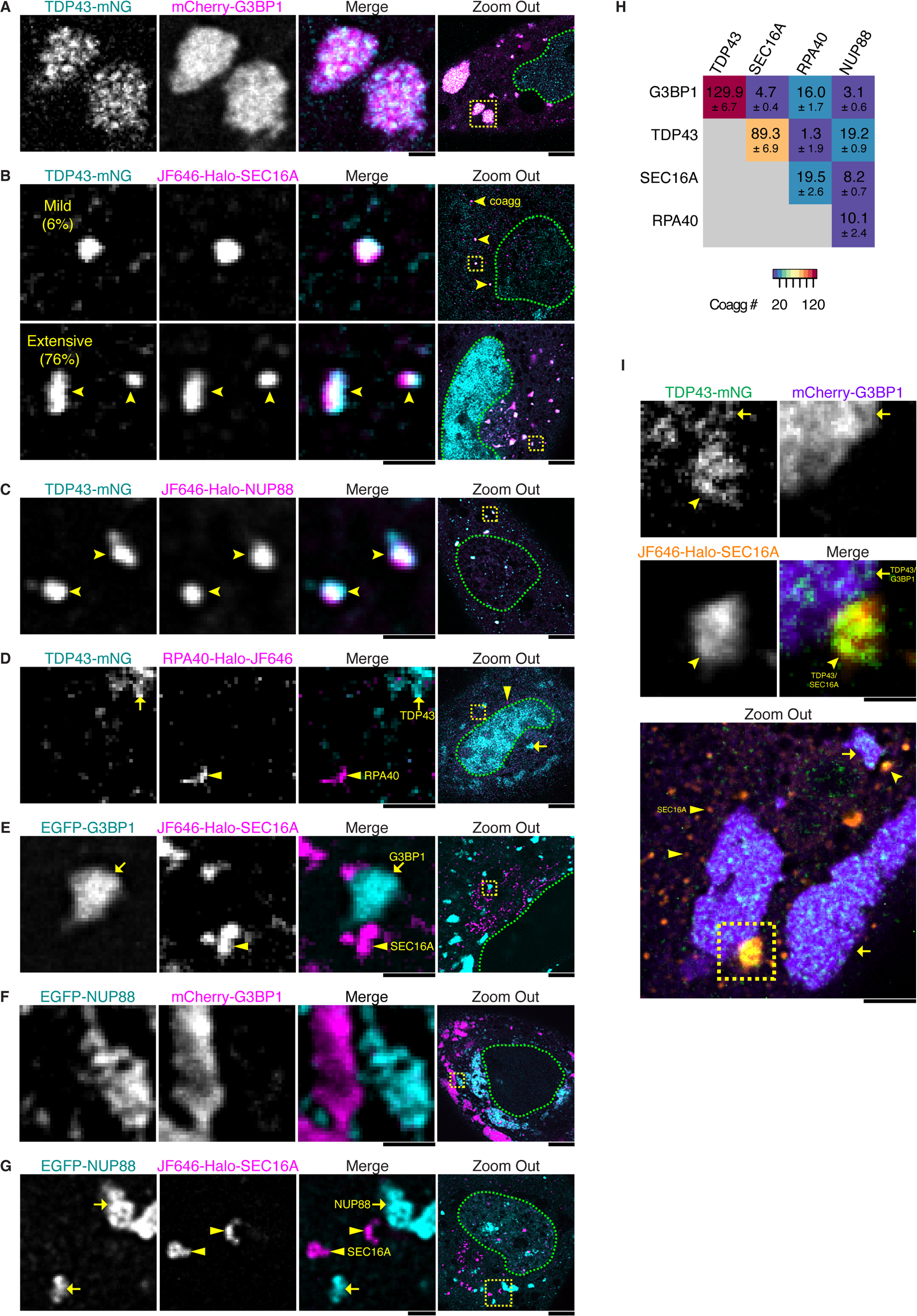
Selective coaggregation among aggregate-forming proteins. (**A**) Representative images showing that TDP43 coaggregated with G3BP1 in stress granule (SG). In “Zoom out”, the yellow box indicates the region shown in zoom-in views, and green dashes demarcate the nucleus. Scale bars except in “Zoom Out” represent 1 μm, and represent 5 μm in “Zoom Out”. (**B**) Representative images for SEC16A inclusions enriching TDP43 mildly (< 10%) and more extensively (≥ 10%) are shown. Arrowheads: coaggregates (coaggs). (**C**) Representative images showing TDP43 coaggregation with NUP88 in the cytoplasm. (**D**) Representative images showing that TDP43 rarely coaggregated with RPA40. Arrows: TDP43 aggregates; triangles: RPA40 aggregates. (**E**) Representative images showing that G3BP1 rarely coaggregated with SEC16A. Arrows: G3BP1 aggregates; triangles: SEC16A aggregates. (**F**) Representative images showing that G3BP1 rarely coaggregated with NUP88. (**G**) Representative images showing that NUP88 rarely coaggregated with SEC16A. Arrows: NUP88 aggregates; triangles: SEC16A aggregates. (**H**) Coaggregation matrix of TDP43, G3BP1 and newly identified aggregation-prone proteins SEC16A, RPA40 and NUP88. The number in each colored square is the mean number of coaggs per cell ± standard deviation (s.d.) derived from analysis of the samples corresponding to (A - G) and Figure S3A. N ≥ 3 x 15. (**I**) Representative images showing TDP43/SEC16A, TDP43/G3BP1 coaggs (*i.e.* TDP43-containing SGs) and SEC16A inclusions without TDP43 in the same cell, which are marked by arrowheads, arrows and triangles, respectively.

Notably, G3BP1, SEC16A and NUP88 rarely coaggregated with each other even though they all showed some degree of coaggregation with TDP43 (Figure 2E - H). We further confirmed that the coaggs of TDP43/G3BP1 (*i.e.* TDP43-containing SGs) and TDP43/SEC16A are orthogonal types of aggregates by simultaneously labelling TDP43, G3BP1 and SEC16A with different fluorescent tags. Even though TDP43/G3BP1 and TDP43/SEC16A coaggs can co-exist in the same cells and some appear adjacent to each other, they do not fuse or colocalize (Figure 2I). Importantly, RNA fluorescence in situ hybridization (FISH) revealed that TDP43/G3BP1 coaggs contained abundant poly-adenine (polyA) RNAs, as previously reported (Jain et al., 2016), whereas TDP43/SEC16A coaggs were devoid of polyA RNA (Figure S4A).

Timelapse imaging revealed that TDP43 partitions dynamically between SGs and TDP43/SEC16A coaggs. In about half of the cells that formed SGs under heat stress, SGs were observed to undergo spontaneous dissolution to below 10% of the initial volume. In these cells, TDP43/SEC16A coaggs gradually emerged as SGs were disappearing (Figure S4D, E and Video S1), and exhibited a significantly higher increase of TDP43 intensity in coaggs with SEC16A when compared with cells that maintained SGs (Figure S4F). To test if the partition of TDP43 can be modulated by the level of SGs, we overexpressed G3BP1 or used RNA interference (RNAi) to inhibit G3BP1/2 expression, which respectively stimulated or suppressed SG assembly (Figure S4G and H) (Matsuki et al., 2013, Tourrière et al., 2023). Correspondingly, the percentage of SEC16A inclusions accumulating TDP43 was respectively reduced or increased (Figure S4I). Furthermore, TDP43^5FL^, an RNA-binding-deficient mutant with reduced recruitment to SGs (Mann et al., 2019), displayed increased coaggregation with SEC16A when compared with wild-type (WT) TDP43 (Figure S4B and C). Together, these results suggest that TDP43 coaggregation with SEC16A is modulated by the capability of SGs to recruit TDP43.

### TDP43/SEC16A coaggs are formed by TDP43 coalescence with ERES under proteotoxic stress

Whereas TDP43/G3BP1 and TDP43/NUP88 coaggs had been reported previously (Mann et al., 2019, Chou et al., 2018), TDP43/SEC16A coaggs represents a new class of TDP43-containing aggregates, which we further investigated. Immunofluorescence staining revealed the coaggregation of endogenous untagged TDP43 and SEC16A under heat stress or combined with MG132 (Figure S3C and D). TDP43/SEC16A coaggs could also be observed in different cell types including HEK293T (Figure S3E) and motor neurons (elaborated further below). Apart from heating, TDP43/SEC16A coaggs could be induced by other proteotoxic stresses such as treatment with MG132 or the autophagy inhibitor Bafilomycin A1 (Baf) (Figure 3A and B). Recently, it was reported that in certain types of cells treated with elevated concentration of sodium chloride (NaCl) for 4 hr, ERES become enlarged liquid-like condensates, termed Sec bodies (SBs) (Zacharogianni et al., 2014, van Leeuwen et al., 2022), but whether SBs contain TDP43 was unknown. We thus treated RPE-1 cells with the same stress, but it did not induce TDP43/SEC16A coaggregation (Figure 3A and B), indicating that TDP43/SEC16A coaggs were distinct from SBs.

**Figure 3.**
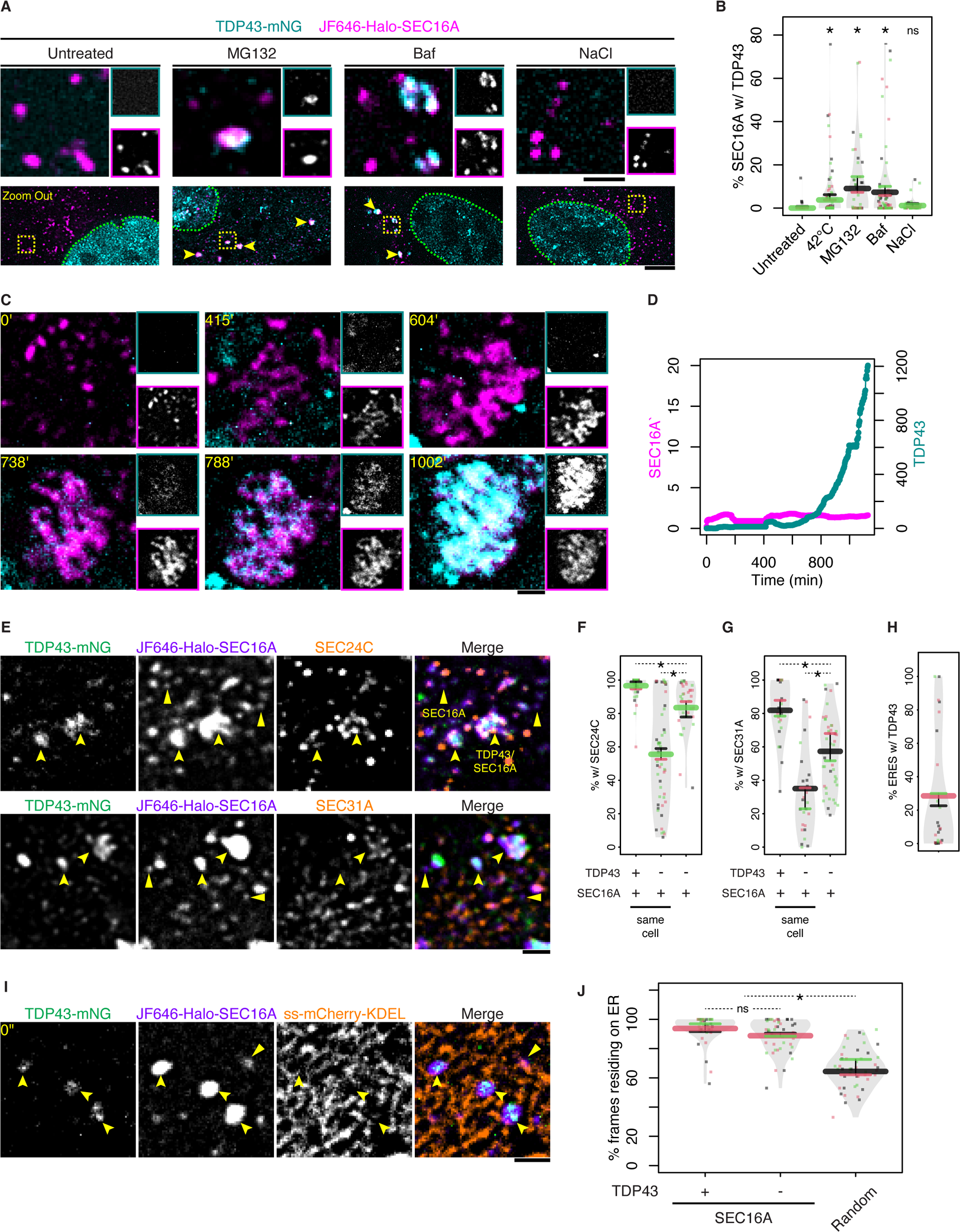
Proteotoxic stress induces TDP43 aggregation with preformed ERES. (**A**) Representative images showing TDP43/SEC16A coaggs in cells treated with 10 μM MG132 or 100 nM Bafilomycin A1 (Baf) for 12 - 16 hr. In comparison, untreated cells or cells treated with 200 mM NaCl for 4 hr were shown. Cyan box: TDP43-mNG channel; magenta box: JF646-Halo-SEC16A channel. In “Zoom Out”, arrowheads indicate coaggs, yellow boxes indicate regions shown in zoom-in views, and green dashes demarcate the nuclei. Scale bars except in “Zoom Out” represent 1 μm, and represent 5 μm in “Zoom Out”. (**B**) Quantification of the experiments as in (A) and Figure 2B showing the volume percentage (% [v/v]) of SEC16A inclusions that enrich TDP43 under the indicated conditions. N ≥ 3 x 15. Asterisks and “ns” respectively stand for significant (p-value ≤ 0.05) and non-significant (p-value > 0.05) in t-tests between untreated and each other condition. (**C**) Selected frames from a representative timelapse recording of ERES in a cell shifted to 42°C. Imaging started at 60 min after the temperature shift. (**D**) Quantification of the experiment in (C) showing the relative intensities of SEC16A and TDP43 over time in ERES (normalized to their respective initial values). (**E**) Representative images of SEC24C or SEC31A immunostaining in cells with TDP43/SEC16A coaggs. Arrowheads: coaggs; triangles: SEC16A inclusions without TDP43. (**F and G**) Quantification of the experiments as in (E) showing the percentage of TDP43/SEC16A coaggs that contained SEC24C or SEC31A, compared with that of SEC16A inclusions without TDP43 in the same cells or in cells without TDP43/SEC16A coaggs. N ≥ 3 x 20. (**H**) Quantification of the experiments as in (E) showing that the volume percentage of fully formed ERES (defined as SEC16A/SEC31A inclusions) that contained TDP43 in cells with TDP43-ERES. N ≥ 3 x 20. (**I**) Representative images of the ER-localization of TDP43/SEC16A coaggs (indicated by arrowheads) and SEC16A inclusions without TDP43 (indicated by triangles). The timelapse recording can be found in Video S3. (**J**) Quantification of the experiments as in (I) and Video S3 showing the percentage of frames in which all TDP43/SEC16A coaggs or all SEC16A inclusions without TDP43 were associated with the ER. As a control, the percentage of ER-residing frames of simulated, randomly distributed particles of the same number and sizes as TDP43/SEC16A coaggs is shown. N ≥ 3 x 15.

To observe how TDP43/SEC16A coaggs initiate under proteotoxic stress, we performed timelapse imaging of cells subject to heat stress at 42°C. Rather than co-assembly with soluble SEC16A to produce coaggs *de novo*, TDP43 gradually aggregated in pre-existing SEC16A inclusions (Figure S5A, 3C, D and Video S2). These SEC16A inclusions often became enlarged and/or clumped as TDP43 accumulated (Figure 3C and S5B) and can be observed under various stress conditions (Figure 3A). The average size of TDP43/SEC16A coaggs, as estimated by structured illumination microscopy (SIM), was ∼ 0.25 μm^3^, substantially larger than that of SEC16A inclusions without TDP43 (∼ 0.06 μm^3^) but smaller than SGs containing TDP43 (∼ 2.5 μm^3^) (Figure S5C). Nonetheless, ∼ 20% TDP43/SEC16A coaggs had similar size as SEC16A inclusions without TDP43, indicating that size alone was insufficient to distinguish between them (Figure S5D).

Then we examined whether TDP43/SEC16A coaggs contained the COPII protein complex, which consists of SEC23/SEC24 subcomplex as the inner layer and SEC13/SEC31 as the outer layer, and are recruited to SEC16A inclusions to form mature ERES (Figure S5A) (Kurokawa and Nakano, 2019, Budnik and Stephens, 2009, Saito et al., 2017). Notably, not all SEC16A inclusions colocalized with SEC24C or SEC31, but almost all TDP43/SEC16A coaggs contained both COPII proteins and the percent COPII recruitment is much higher than that of SEC16A inclusions without TDP43 (Figure 3E - G), indicating that TDP43/SEC16A coaggs originated from mature ERES. In cells that contained TDP43/SEC16A coaggs, the volume percentage [v/v] of (mature) ERES (defined as SEC16A/SEC31A foci) that enrich TDP43 is ∼ 30% on average (Figure 3H). Interestingly, in these cells, SEC16A inclusions without TDP43 exhibited further reduced levels of COPII recruitment compared to those in cells free of TDP43/SEC16A coaggs (Figure 3F and G), suggesting that TDP43/SEC16A coaggs sequestrate COPII subunits away from other SEC16A inclusions.

Further supporting that TDP43/SEC16A coaggs are resulted from TDP43 aggregation with ERES, timelapse imaging revealed that TDP43/SEC16A coaggs remained associated with the ER like other SEC16A inclusions (Figure 3I, J and Video S3). Therefore, we hereafter refer to TDP43/SEC16A coaggs as “TDP43-ERES”, and ERES without TDP43 aggregation as “ordinary ERES”.

Because TDP43 is an RNA-binding protein but TDP43-ERES did not contain polyA RNA (Figure S4A), we suspected that nascent or unfolded TDP43 were enriched in ERES. To test this, we employed a system of Halo-tagged TDP43 and different HaloTag ligand dyes to differentiate between pre-existing (mature) versus newly synthesized (new) TDP43 (Figure S5E). To facilitate this analysis, we used MG132 to inhibit the proteasomal degradation of soluble TDP43 (Scotter et al., 2014). Although both the mature and new pools of TDP43 aggregated in ERES (Figure S5F and Video S4), the ratio between the new and mature TDP43 in ERES, normalized to the new/mature ratio of TDP43 in the whole cell, was always greater than one (Figure S5G and H), indicating that nascent rather than mature TDP43 was the favored substrate for aggregation in ERES. Such selectivity for nascent TDP43 was not attributed to different dye properties, as the preference was not abolished by switching the use of dyes (Figure S5I). Interestingly, the normalized new/mature ratio gradually decreased during the lifetime of TDP43-ERES (Figure S5G and H), suggesting that nascent TDP43 formed the seed for aggregation, to which pre-existing TDP43 was added during aggregate growth. In support of this possibility, the protein translation inhibitor cycloheximide (CHX) strongly suppressed TDP43 aggregation in ERES (Figure S5J and K).

### TDP43-ERES are solid aggregates distinct from SGs or ordinary ERES

In contrast to their highly dynamic nature in SGs or ERES without TDP43 (Mann et al., 2019, Watson et al., 2006), molecules of both TDP43 and SEC16A are immobile in TDP43-ERES. In fluorescence recovery after photobleaching (FRAP) experiments, FP-tagged TDP43 and SEC16A in TDP43-ERES were partially photobleached, but the bleached and unbleached regions hardly mixed over time (Figure 4A, B and Video S5). Quantification of the FRAP data yielded the mobile fractions of TDP43 and SEC16A molecules in coaggs to be roughly 15% and 10%, respectively (Figure 4C). By contrast, TDP43 and G3BP1 in SGs rapidly and fully mixed after partial photobleaching (Figure S6A, B and Video S6) with mobile fractions of each protein to be above 60% (Figure 4C).

**Figure 4.**
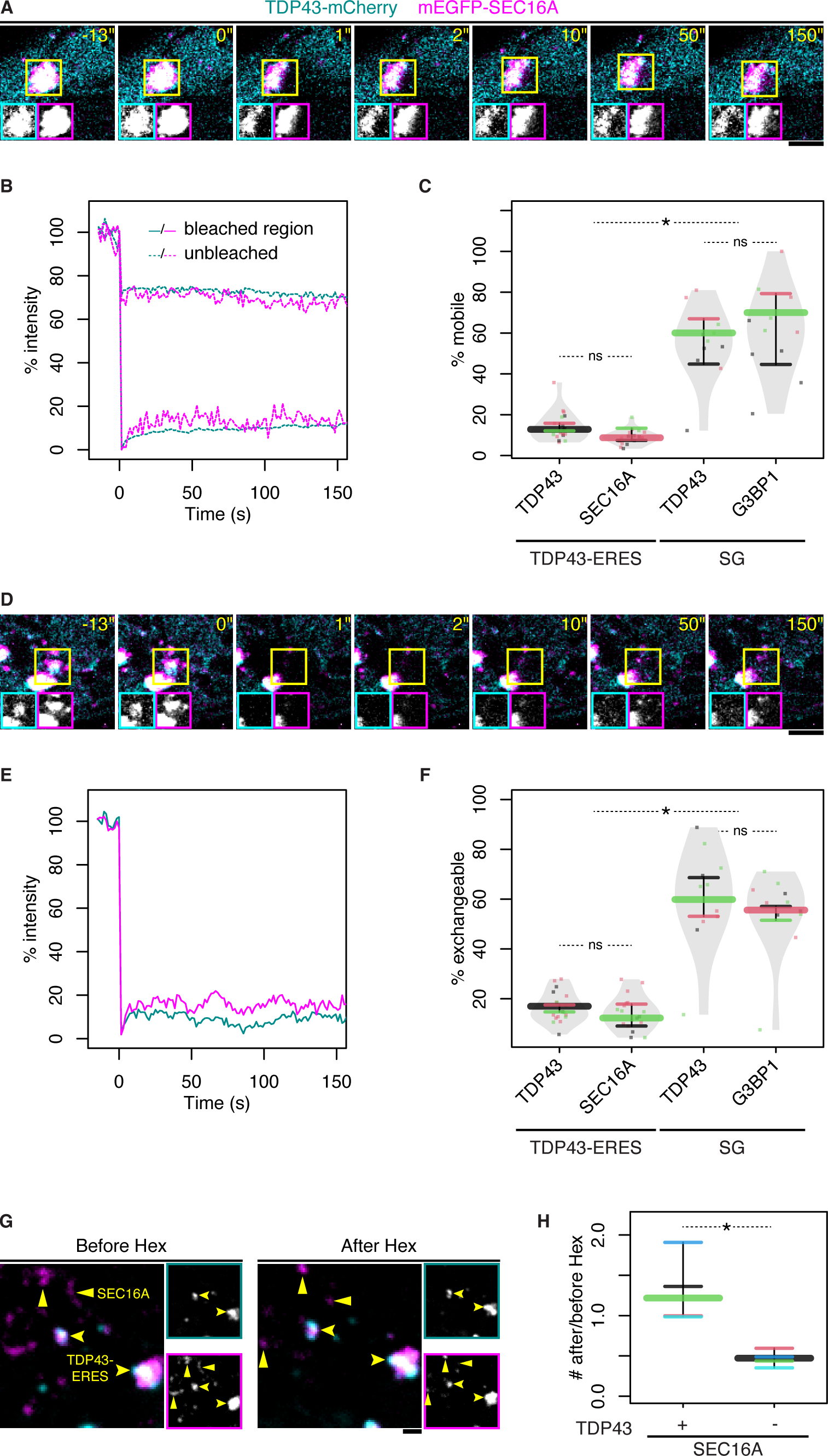
TDP43-ERES are solid-like aggregates. (**A**) Selected frames from a representative timelapse recording of a TDP43-ERES coagg after partial photobleaching. Insets show individual channels of the yellow boxed region. Each scale bar represents 1 μm. (**B**) Quantification of the experiment in (A) showing the relative intensities of TDP43 (cyan lines) and SEC16A (magenta lines) in bleached (solid lines) and unbleached regions (dashed lines) over time. (**C**) Quantification of partial-FRAP experiments as in (A) and Figure S6A comparing the mobile fractions of proteins in TDP43-ERES versus SGs. Three experiments were performed and the average of 3 - 9 aggregates in each experiment was shown as a datapoint. (**D**) Selected frames from a representative timelapse recording of a TDP43-ERES after photobleaching in its entirety. The yellow box marks the position of the coagg immediately before photobleaching. (**E**) Quantification of the experiment in (D) showing the relative intensities of TDP43 (cyan line) and SEC16A (magenta line) over time. (**F**) Quantification of full-FRAP experiments as in (D) and Figure S6C comparing the exchangeable fraction of proteins in TDP43-ERES versus SGs. 3 experiments were performed and the average of 3 - 9 aggregates in each experiment was shown as a datapoint. (**G**) Representative images showing TDP43-ERES (indicated by arrowheads) and SEC16A inclusions without TDP43 (indicated by triangles) in a cell before and after 3.5% [v/v] 1,6-hexanediol (Hex) treatment for 15 min. Insets show individual TDP43-mCherry (cyan) and mEGFP-SEC16A (magenta) channels. (**H**) Quantification of the experiments as in (G) comparing the after/before ratios of TDP43-ERES and SEC16A inclusions without TDP43. N ≥ 3 x 10 and the median values of cells in the same experiments are plotted as horizontal segments.

The solid-like nature of TDP43-ERES was also reflected by a lack of exchange with the cytosol: when coaggregated TDP43 and SEC16A were photobleached entirely, the fluorescence recovery for either protein was below 20% over 2.5 min (Figure 4D - F and Video S7). On the other hand, TDP43 and G3BP1 in SGs rapidly exchanged with the cytosol (Figure S6C - E and Video S8) with more than 55% exchangeable with the cytosol (Figure 4F).

Lastly, TDP43-ERES were resistant to 1,6-hexanediol (Hex) (Figure 4G and H), which disrupts weakly aggregated macromolecules (Düster et al., 2021). In fact, Hex treatment led to a ∼ 30% increase in the number of TDP43-ERES due to the formation of new TDP43/SEC16A coaggs in some cells (Figure 4H and Video S9). By contrast, after Hex treatment for 15 min, ∼ 50% of SEC16A inclusions without TDP43 regardless of size and nearly all SGs dissolved (Figure 4G, H and S6F – H), consistent with previous reports (Wheeler et al., 2016, Zacharogianni et al., 2014, van Leeuwen et al., 2022). Together, the above findings demonstrated that TDP43-ERES were solid aggregates distinct from SGs and ordinary ERES.

### TDP43 aggregation with ERES impairs ER-to-Golgi transport

As ERES is critical for vesicular transport from ER to Golgi, we examined whether TDP43-ERES aggregation compromises its function. Retention Using Selective Hooks (RUSH) is a method in which secretory substrates, such as ManII or TNFα, fused with streptavidin-binding protein (SBP), can be artificially retained in the ER by interaction with the streptavidin anchor. Addition of biotin then releases the retained substrates to enable quantitative analysis of protein trafficking (Figure S7A) (Boncompain et al., 2012). We introduced the RUSH system into cells expressing TDP43-mNG and Halo-SEC16A and assayed the ER-to-Golgi transport of ManII-SBP-mCherry (RUSH-ManII) under heat stress. As expected, in the absence of biotin, RUSH-ManII was retained in the ER and not delivered to Golgi, regardless of whether TDP43 aggregated in ERES (Figure S7B). After incubation with biotin for 1 hr, RUSH-ManII appeared in Golgi in most cells without TDP43-ERES and was no longer seen in the ER (Figure 5A, B and S7C). By contrast, in cells with extensive TDP43-ERES aggregation (> 10% [v/v] SEC16A inclusions with TDP43), a considerable fraction of RUSH-ManII remained in the ER and outside Golgi (Figure 5A and B). Furthermore, overexpression of TDP43-mNG but not mNG alone significantly increased the fraction of cells that displayed ER export defect (Figure 5C), supporting that elevated TDP43 impaired ER-to-Golgi transport.

**Figure 5.**
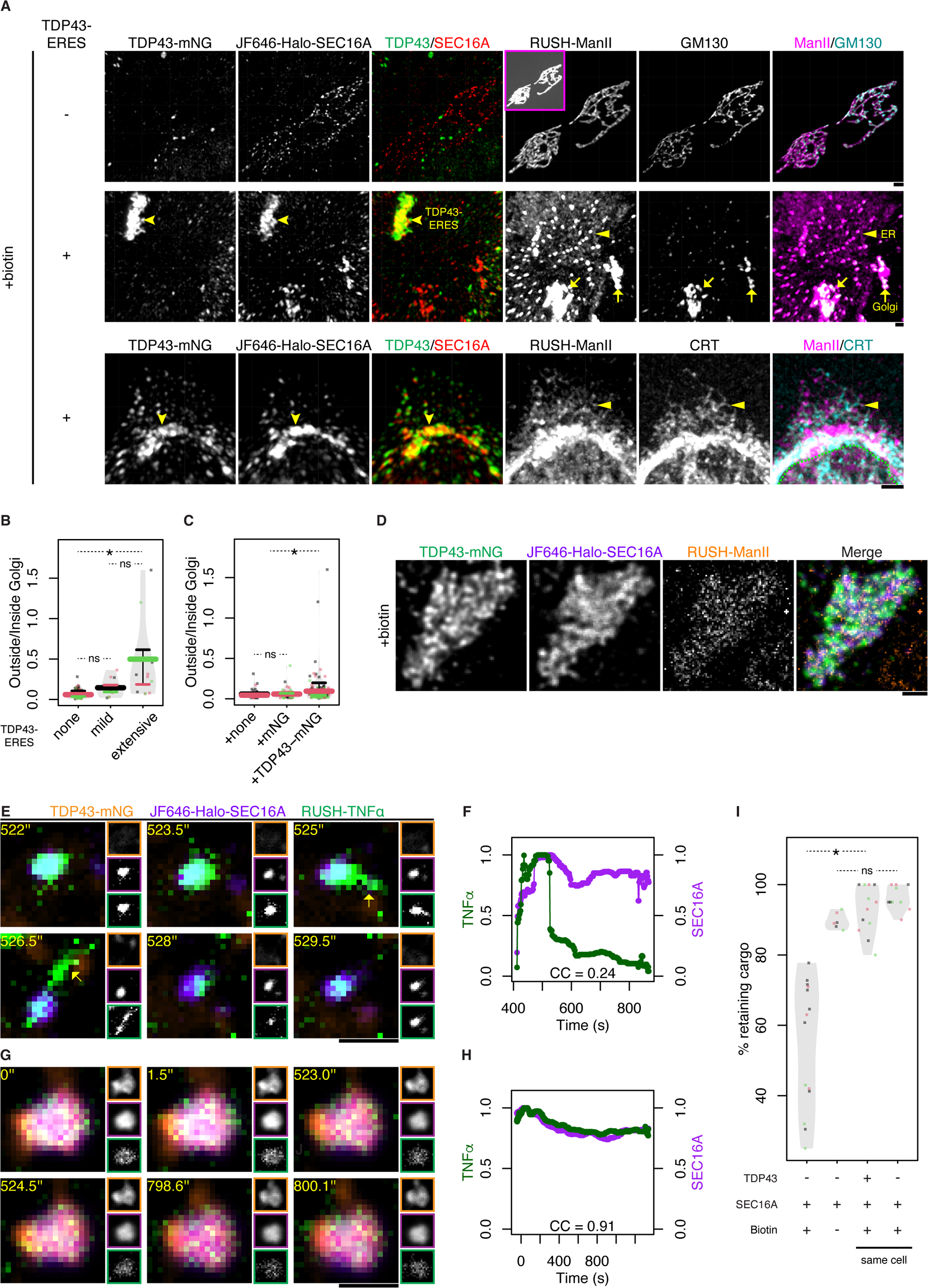
TDP43-ERES aggregation impairs ER-to-Golgi transport. (**A**) Representative images of RUSH assays in cells without (Row1) or with TDP43 coaggregation with ERES (Row 2 and 3). After biotin addition for 1 hr, the subcellular localization of ManII-SBP-mCherry (RUSH-ManII) was imaged with GM130 (Golgi marker) or calreticulin (CRT – ER marker) visualized by using immunofluorescence staining. In Row1, the inset in the RUSH-ManII channel shows a contrast-adjusted view. In Row 2 and 3, arrowheads indicate TDP43-ERES; arrows point to fragmented Golgi; triangles point to RUSH-ManII retention in the ER; green dashes demarcate the nucleus. The 3D projections (view from top) of these cells are displayed. Each scale bar represents 1 μm. (**B**) Quantification of the experiments as in (A) and Figure S7C showing the ratios between RUSH-ManII intensity outside and inside Golgi. Cells were categorized based on their volume percentage of TDP43-ERES (none: 0%; mild: 0 – 10%; severe: ≥ 10%). N ≥ 3 x 15. (**C**) Quantification of cells transfected with RUSH-ManII and Halo-SEC16A only or additionally with mNG or TDP43-mNG showing the outside/inside Golgi ratios of RUSH-ManII intensity. N ≥ 3 x 15. (**D**) Representative images of TDP43-ERES trapping RUSH-ManII after biotin addition for 1 hr. A representative slice of the z-stack is displayed. (**E**) Selected frames of a representative timelapse recording of an ordinary ERES (in cells without TDP43-ERES) during RUSH-TNFα transport from ER to Golgi. Biotin was added at 0’’. Arrows indicate the budding of a tubular transport intermediate. (**F**) Quantification of relative RUSH-TNFα intensity over time (normalized to the max intensity),compared to the relative intensity of SEC16A in the ERES tracked in (E). CC: cross-correlation between RUSH-TNFα intensity and SEC16A intensity. (**G**) Selected frames of a representative timelapse recording of a TDP43-ERES during RUSH-TNFα assay. (**H**) Quantification of RUSH-TNFα and SEC16A intensities over time (normalized to their respective max intensities) in the TDP43-ERES tracked in (G). (**I**) Quantification of the percentage of TDP43-ERES and ERES without TDP43 in either the same or different cells that retained RUSH-TNFα in the experiments as in (E - H) and Figure S8D and E. The criterion for cargo retention is CC ≥ 0.5. N ≥ 3 x 3. Cells without biotin supplement (N = 3 x 2) were included as a control.

Notably, RUSH-ManII were retained in TDP43-ERES after biotin addition for 1 hr (Figure 5D), suggesting that TDP43 aggregation caused ERES dysfunction to trap secretory cargos recruited to these abnormal sites. To directly test this hypothesis, we took advantage of another RUSH reporter, TNFα-SBP-mCherry (RUSH-TNFα), which is pre-enriched in ERES prior to biotin supplement (Weigel et al., 2021). After adding biotin, ordinary ERES that already accumulated RUSH-TNFα efficiently exported this substrate via characteristic tubular transport intermediates/vessels (Figure 5E), consistent with previous observations (Weigel et al., 2021). As such, the intensity of RUSH-TNFα in ordinary ERES decreased dramatically after cargo release while that of SEC16A remained relatively stable (Figure 5F). By contrast, TDP43-ERES, although similarly recruited RUSH-TNFα (Figure S8A and B), failed to form tubular transport intermediates to release this cargo, such that the intensity of RUSH-TNFα remained stable and/or correlated with SEC16A intensity over time - the cross-correlation (CC) between RUSH-TNFα and SEC16A intensities was close to 1 (Figure 5G and H). We hence calculated the CC between RUSH-TNFα and SEC16A intensities for TDP43-ERES and ordinary ERES in our timelapse recordings and defined CC ≥ 0.5 as the benchmark for cargo retention. Based on this criterion, the percentage of TDP43-ERES that retained RUSH-TNFα was significantly higher than that of ordinary ERES (Figure 5I). Surprisingly, in cells with TDP43-ERES, a lower percentage of SEC16A inclusions without TDP43 enriched RUSH-TNFα (Figure S8A and B) and were more likely to fail in cargo release (Figure S8D, E and 5I), suggesting that TDP43-ERES had dominant effects over ordinary ERES in the same cells.

### Artificially induced TDP43 aggregation at ERES inhibits ER-to-Golgi transport

To further investigate whether TDP43-ERES aggregation, rather than cytosolic TDP43 or any pleiotropic effects of heat stress, was the cause of ERES dysfunction, we employed a method to directly aggregate TDP43 in ERES. When TDP43 and SEC16A were respectively fused with three copies of FKBP and FKBP-rapamycin binding domain bearing T2098L mutation (FRB*), the rapamycin analog AP21967 (AP), which dimerizes FKBP and FRB*, directly induced TDP43/SEC16A coaggregation. As a control, cells expressing TDP43 and SEC16A without FKBP or FRB* tagging were not responsive to AP (Figure S9A, 6A and B). Furthermore, like the TDP43-ERES induced by proteotoxic stresses, AP-induced TDP43/SEC16A coaggs contained COPII proteins (Figure 6C), and 3xFKBP-TDP43-mNG and 3xFRB*-Halo-SEC16A molecules inside were immobile as revealed by FRAP (Figure 6D).

**Figure 6.**
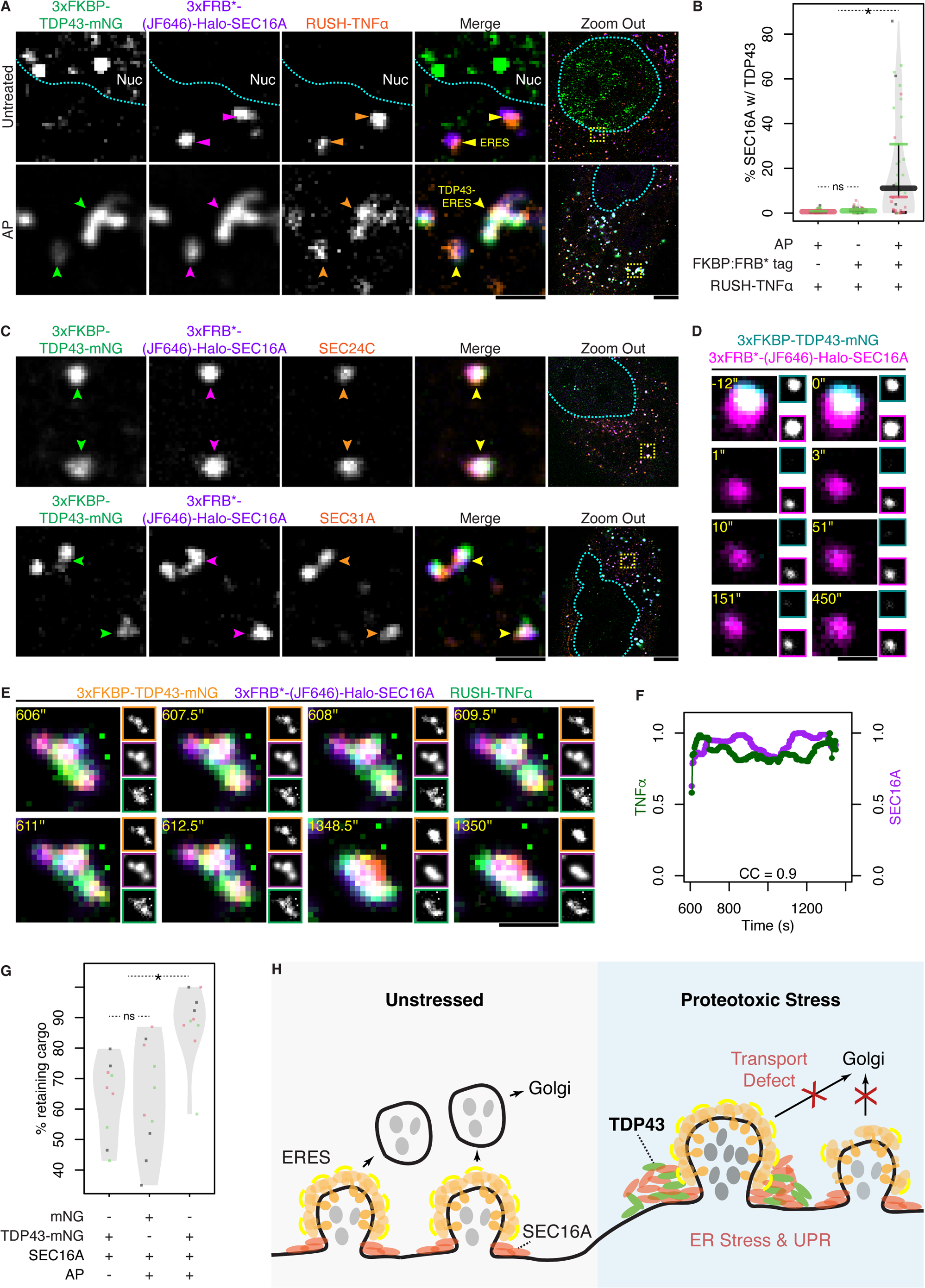
Artificially induced TDP43-ERES retain secretory cargo. (**A**) Representative images of cells cultured in biotin-free media expressing 3xFKBP-TDP43-mNG, 3xFRB*-Halo-SEC16A and RUSH-TNFα without or with AP21967 (AP) treatment for 6 - 10 hr. Arrowheads and triangles indicate TDP43-ERES and ordinary ERES, respectively. In “Zoom out”, yellow boxes mark the regions displayed in zoom-in views, and cyan dashes demarcate the nuclei. Scale bars except in “Zoom Out” represent 1 μm, and represent 5 μm in “Zoom Out”. (**B**) Quantification of the experiments as in (A) and Figure S9B showing the volume percentage of SEC16A inclusions that enriched TDP43 under the indicated conditions. N ≥ 3 x 15. (**C**) Representative images of SEC24C or SEC31A immunostaining in cells with AP-induced TDP43/SEC16A coaggs. (**D**) Selected frames from a representative timelapse recording of an AP-induced TDP43-ERES after photobleaching. Insets show individual channels. 3xFKBP-TDP43-mNG was entirely photobleached whereas 3xFRB*-(JF646)-Halo-SEC16A was partially bleached. The scale bar represents 0.5 μm. (**E**) Selected frames of a representative timelapse recording of an AP-induced TDP43-ERES during RUSH-TNFα assay. (**F**) Quantification of RUSH-TNFα and SEC16A intensities over time (normalized to their respective max intensities) in the TDP43-ERES tracked in (E). (**G**) Quantification of the percentage of AP-induced TDP43(-mNG)-ERES, mNG-ERES and ERES without TDP43 (in cells without AP treatment) that retained RUSH-TNFα in the experiments as in (E) and Figure S9C - F. N ≥ 3 x 3. (**G**) Working model of TDP43 aggregation at ERES causing retention of transport cargos. TDP43-ERES also exert a dominant effect over ordinary ERES in the same cells possibly through sequestration of essential transport factors. The ER-to-Golgi transport defect caused by TDP43-ERES coaggregation may further induce ER stress.

Next, we examined whether AP-induced TDP43 aggregation at ERES affected cargo export from the ER. The AP-induced TDP43-ERES still enriched RUSH-TNFα (Figure 6A), but this cargo was not exported after biotin addition (Figure 6E and F). By contrast, in the absence of AP, cargo release from ERES was unaffected, suggesting that FKBP or FRB tagging alone did not compromise ERES function (Figure S9C, D and 6G). As an additional control, AP-induced recruitment of 3xFKBP-mNG to ERES but did not cause ERES to retain RUSH-TNFα (Figure S9E, F and 6G). These results suggest that TDP43 aggregation in ERES is sufficient to impair ER-to-Golgi transport (Figure 6H).

### TDP43-ERES are found in motor neurons of ALS patients

It was recently proposed that the diffuse punctate cytoplasmic staining (DPCS) of TDP43 are aggregates on the ER membrane and may serve as the earliest marker of ALS pathology (Kon et al., 2022). In addition, motor neurons derived from the human induced pluripotent stem cells (hiPSCs) of ALS patient or found in their post-mortem sections exhibit attenuated vesicle transport, significant ER stress and activation of Unfolded Protein Response (UPR), which contribute to neuronal dysfunction and death (Ilieva et al., 2007, Oyanagi et al., 2008, Matus et al., 2013, Medinas et al., 2018, Medinas et al., 2019, Nishitoh et al., 2008). We therefore tested if TDP43 aggregation in ERES is elevated in ALS patient-derived motor neurons. *C9ORF72* is the most mutated gene in ALS patients (van Rheenen et al., 2021, Balendra and Isaacs, 2018). We induced motor neurons (iMNs) from human embryonic stem cells (hESCs) or hiPSCs derived from ALS patients bearing *c9orf72* (*c9*) mutations (NDS00268, NDS00269 and NDS00270) and non-ALS (control) individuals (BJ, GM23720 and H9) (Figure S10A and B), and stained TDP43 and SEC16A by immunofluorescence. ALS patient-derived iMNs displayed significantly increased level (volume percentage) of TDP43-ERES (Figure 7A and B).

**Figure 7.**
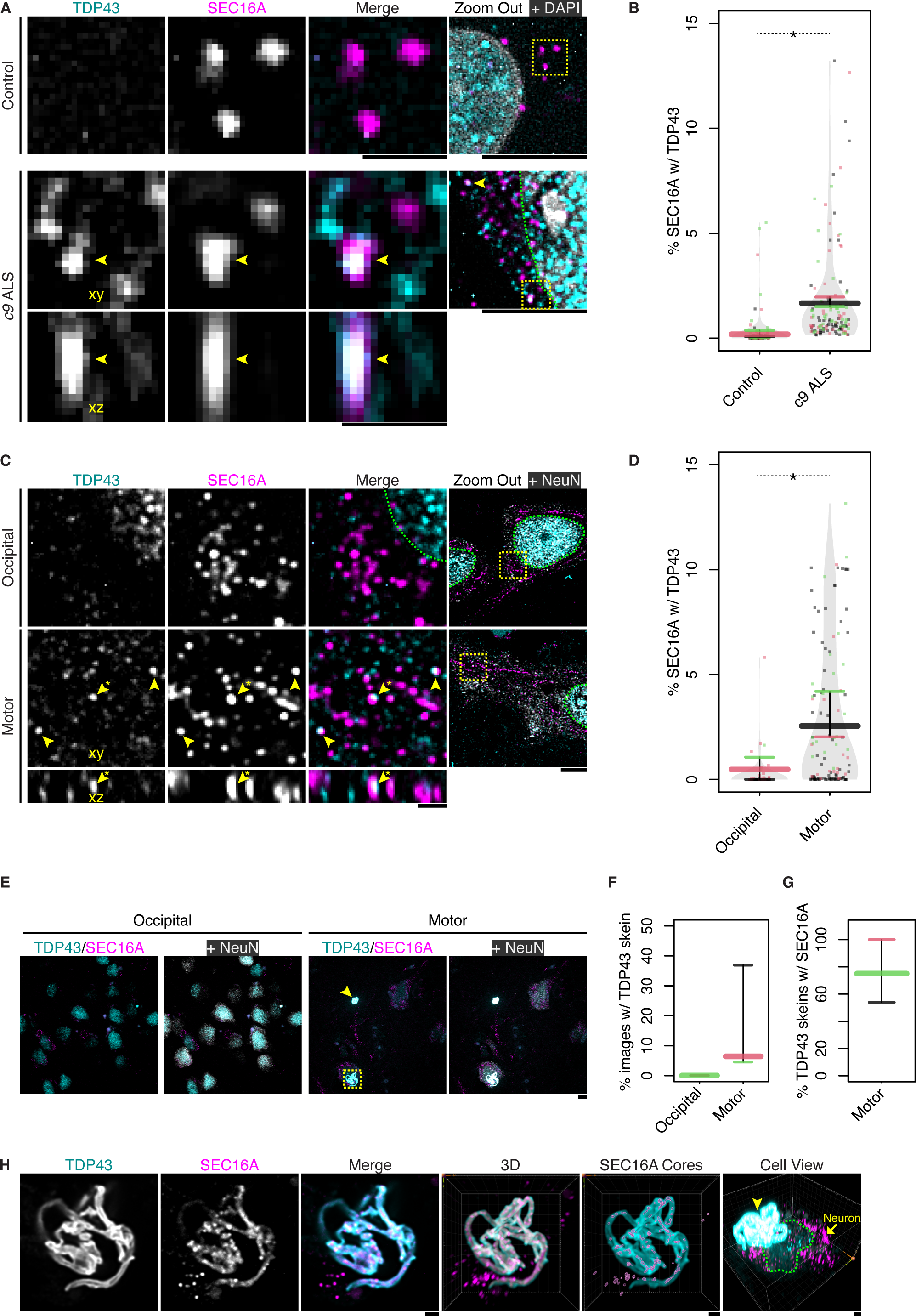
Coaggregation of TDP43 with ERES in ALS-affected motor neurons. (**A**) Representative images of immunofluorescence staining of TDP43 and SEC16A in MNs induced from a non-ALS (Control) hiPSC line (BJ) and an ALS patient hiPSC line (NDS00270) that carries the *c9orf72* mutation (*c9*). In “Zoom out”, yellow boxes mark the regions displayed in zoom-in views, and green dashes demarcate the nuclei. The sectioning of the TDP43/SEC16A coagg along the **xz** plane is displayed beneath the **xy** view. Scale bars except in “Zoom Out” represent 1 μm, and represent 5 μm in “Zoom Out”. (**B**) Quantification of the experiments as in (A) showing the volume percentage of SEC16A inclusions that contained TDP43. 3 normal plus 3 patient hESC/hiPSC lines were used, and 30 – 50 iMNs derived from each cell line (coded with different colors) were imaged (*i.e.* N ≥ 3 x 30). (**C**) Representative images of the immunofluorescence staining of TDP43, SEC16A and the neuronal marker NeuN (RBFOX3) in the post-mortem sections of the occipital cortex and motor cortex from the same sALS patient. The arrowheads with asterisk indicate the same TDP43/SEC16A coagg in **xy** and **xz** views. (**D**) Quantification of the experiments as in (C) showing the volume percentage of SEC16A inclusions containing TDP43 in the occipital cortex versus motor cortex of 3 different sALS patients (color-coded). At least 30 images were taken for each patient (*i.e.* N ≥ 3 x 30). (**E**) Representative images of TDP43, SEC16A and NeuN immunostaining in the occipital cortex and motor cortex of the same sALS patient. 2 skein-like TDP43/SEC16A aggregates in the motor cortex are boxed and indicated by arrowhead, with their respective zoom-in views displayed in (H) and Figure S10E. The max intensity projection of confocal z-stacks are shown. Each scale bar represents 5 μm. (**F**) Quantification of the percentage of confocal z-stack images containing skein-like TDP43 aggregates in the occipital versus motor cortex of 3 different sALS patients (color-coded) in the experiments as in (E). N ≥ 3 x 30. (**G**) Quantification of the percentage of TDP43 skeins that contained SEC16A in the motor cortex of sALS patients in the experiments as in (E). At least 5 TDP43 skeins were found in each sALS patient (*i.e.* N ≥ 3 x 5). (**H**) Zoom-in views of the TDP43/SEC16A skein-like aggregate boxed in (E). “SEC16A Cores” shows the contours of high SEC16A fluorescence intensity; “Cell View” shows a neuron (indicated by the arrow) beneath the TDP43/SEC16A skein (indicated by the arrowhead), and the nucleus of this neuron is demarcated by green dashes.

About 90% of ALS cases are characterized as sporadic ALS (sALS) (Masrori and Van Damme, 2020), so we used immunofluorescence staining to analyze TDP43/SEC16A coaggregation in the post-mortem brain sections of sALS patients. In neurons of the (ALS-affected) motor cortex but not the (unaffected) occipital cortex, TDP43 fibrils could be observed in the cytoplasm while TDP43 were depleted from the nucleus (Figure S10C and D), as previously reported (Neumann et al., 2006, Suk and Rousseaux, 2020). In the motor cortex, TDP43-ERES were found at higher level (volume percentage) compared with the occipital cortex (Figure 7C and D), and the morphology and subcellular distribution of TDP43-ERES are similar to those of DCPS, *i.e.* both are diffusive puncta in the cytoplasm (Figure 7C). In addition, we observed skein-like TDP43 aggregates in the extracellular space of the motor cortex but not occipital cortex (Figure 7E and F), and most of these TDP43 skeins contained SEC16A (Figure 7E and G). Notably, along these TDP43/SEC16A skeins, there were high intensity cores of SEC16A (Figure 7H and S10E), which resembled the clustered form of TDP43-ERES observed in heat stressed RPE-1 cells (Figure 3C and S5B). The above observations are consistent with the notion that TDP43 aggregation at ERES plays a role in ALS cellular pathology.

## Discussion

Protein aggregation, or clumping, results in the formation of liquid-like or solid inclusions. Recently, it was found that SGs and nucleoli can undergo phase transition from liquid-like to solid aggregates by accumulating misfolded proteins (Mateju et al., 2017, Patel et al., 2015, Frottin et al., 2019). In the present study, we utilized Hsp104^DWB^-mEGFP-FLAG in an affinity-based approach to identify the protein components of aggregates that contain misfolded proteins in human cells. Among the 337 aggregation-prone proteins identified, there are not only components of SGs and nucleoli but also SEC16A, which was suggested to undergo LLPS to constitute the scaffold of ERES (Watson et al., 2006, Budnik and Stephens, 2009, Gallo et al., 2023). Our data further show that, under heat stress, a fraction of ERES contain misfolded proteins as revealed by Hsp104^DWB^-mEGFP binding, and some accumulate TDP43 and become solid. Therefore, ERES are a new type of precursors for solid aggregate formation. Whereas TDP43-containing SGs can remain liquid-like for hours to days (Mann et al., 2019, Mateju et al., 2017, Patel et al., 2015), ERES turn solid soon after stress- or AP-induced TDP43 aggregation. Importantly, SGs that remain in liquid-like state have been proposed to play certain cytoprotective roles, such as storing mRNAs other than those involved in stress response (Nover et al., 1989, Khong et al., 2017, Hofmann et al., 2021) although SGs can be crucibles for solid aggregate formation during prolonged stress (Mateju et al., 2017, Patel et al., 2015). In this study, we observed that SG formation suppresses TDP43 aggregation at ERES, which might represent another aspect of cytoprotection by SGs worth of further exploration.

Previous study showed that 40 – 60% of TDP43 is synthesized by ER-attached ribosomes by ribosome profiling (Reid and Nicchitta, 2012, Jan et al., 2014), and TDP43 can interact with lipid bilayers through its prion-like domain (Lim et al., 2016), which may contribute to its concentration on the ER surface and recruitment to ERES. In addition, ERES was recently shown to form at highly connected regions of the ER with “high search rate” and consist of interwoven networks of membranes (Weigel et al., 2021), which potentially traps the newly synthesized TDP43. Speculatively, when enriched at ERES, the prion-like C-terminal domain of TDP43 could interact directly with the intrinsically disordered region (IDR) of SEC16A, which spans more than 50% of the latter’s sequence (Prasad et al., 2019, Gallo et al., 2023). Consistent with our previous findings in yeast that stress-induced aggregates emerge from the cytoplasmic surface of the ER and are likely to be seeded by newly synthesized proteins (Zhou et al., 2014), in this study, we found that the nascent TDP43 is more prone to aggregation in ERES than pre-existing TDP43 and blocking protein synthesis by CHX strongly suppressed such aggregation. This may be explained by a combination of the proximity of nascent TDP43 to ERES, the fact that newly synthesized proteins are more prone to stress-induced misfolding and damage (Vabulas and Hartl, 2005, Medicherla and Goldberg, 2008), and possibly a lack of RNA-binding or certain post-translational modifications on nascent TDP43 (Mann et al., 2019).

TDP43-ERES are usually larger in size than ordinary ERES, like the recently reported Sec bodies (SBs), which are enlarged ERES (> 300 nm in radius) retaining liquid-like properties (Zacharogianni et al., 2014, van Leeuwen et al., 2022). However, TDP43-enriched ERES are distinct from SBs in multiple aspects. First, pronounced SBs were found only in *Drosophila melanogaster* (fruit fly) S2 cells and a specific mammalian cell line, the rat insulinoma INS-1, but not in several other cell lines studied (van Leeuwen et al., 2022). By contrast, we observed TDP43-enriched ERES in multiple human cell lines of different tissue origins, including RPE-1, HEK293T and motor neurons. Second, SBs were induced under specific conditions such as high NaCl concentration (van Leeuwen et al., 2022) whereas TDP43-enriched ERES were induced by various proteotoxic stresses but not NaCl supplement. Lastly, SBs are liquid-like structures whereas ERES that assimilated TDP43 were solid. However, the above differences cannot rule out a possible link between SBs and TDP43.

It has been questioned whether protein aggregation is the cause or result of cellular dysfunction and whether aggregation is protective against proteotoxic damages (Kopito, 2000, Soto, 2003, Kayatekin et al., 2014, Bolognesi et al., 2019, Bence et al., 2001, Park et al., 2013, Garcia et al., 2021). Our findings suggest that TDP43 aggregation in ERES is deleterious, as it compromises the function of ERES in cargo export from the ER. Our data also suggest that TDP43-ERES exerts a dominant effect over ordinary ERES, potentially through sequestration of COPII subunits or other limiting factors regulating COPII vesicle release. As such, when TDP43 aggregates in over 10% [v/v] of SEC16A inclusions, the ER-to-Golgi transport is globally delayed. As TDP43-ERES are solid aggregates with limited exchange of molecules with the cytosol, they cannot spontaneously disassemble. Therefore, the inhibitory effect of TDP43-ERES aggregation on the secretory pathway may not be readily reversed after stress attenuation, which would be different from the regulatory inhibition of ER export by reversible SG recruitment of SEC23 and SEC24, but not SEC16A, reported previously (Zappa et al., 2019).

TDP43-ERES are able to recruit cargos but not to generate tubular transport vessels to release them, suggesting that TDP43-ERES are defective in COPII vesicle budding. Such dysfunction could result from the inhibitory effect of TDP43 on the GTPase cycle of COPII complex, or on the mechanics of vesicle budding due to liquid-to-solid phase transition associated with TDP43 aggregation. Recent studies suggest that liquid-like assemblies can promote membrane bending or budding under certain conditions (Yuan et al., 2021, Mangiarotti et al., 2022). Such potential effect of SEC16A condensates on the ER membrane could be disrupted by TDP43-induced phase transition. Moreover, ERES was recently found to generate tubule vessels of connected COPII vesicles rather than separate ones as the transport intermediate (Weigel et al., 2021), and solid aggregates might mechanically block the elongation of these pearled tubules.

Finally, our results show that TDP43 aggregation in ERES occurs in ALS patient hiPSC-induced motor neurons (iMNs) and post-mortem samples of motor cortices. Supporting the disease relevance of our findings, SEC16A mutations were recently identified as a risk factor for ALS (Zhang et al., 2022), and defects in intracellular vesicle transport were reported in ALS patient iMNs (Shi et al., 2018). In addition, ER stress and activation of UPR leading to ER expansion are widely recognized hallmarks of ALS pathology (Ilieva et al., 2007, Oyanagi et al., 2008, Matus et al., 2013, Medinas et al., 2018, Medinas et al., 2019). Previous studies indicated that ER-to-Golgi transport is inhibited by certain TDP43, SOD1, FUS or UBQLN2 mutants (Soo et al., 2015, Halloran et al., 2020), which, however, are rare among ALS patients (van Rheenen et al., 2021, Balendra and Isaacs, 2018). Our findings on the harmful TDP43 aggregation in ERES suggest a more general mechanistic explanation for the clinical observations of ER stress. Given the morphological similarity between intracellular TDP43-ERES coaggregates and the DPCS, this type of aggregates could potentially serve as an early diagnostic marker or therapeutic target for alleviating the toxic effects of TDP43 aggregation in ALS, whereas extracellular TDP43/SEC16A skeins might represent clustered TDP43-ERES aggregates released after cell death.

## Materials and Methods

### Mammalian cell culture and plasmid transfection

Cell culture was performed according to the American Type Culture Collection (ATCC) guidelines with minor modifications. Briefly, RPE-1 cells were maintained in DMEM:F12 (Thermo 11320082) + 10% [v/v] FBS (Thermo 16140071) and passaged using 0.05% [w/v] trypsin solution (Thermo 25300120). HEK293T cells were cultured in DMEM (Thermo 11965118) + 10% [v/v] FBS and passaged similarly. For high-resolution microscope imaging, cells were seeded on glass-bottom dishes (Iwaki 3930-035 or 3931-035) pre-coated with 10 μg/mL fibronectin (Sigma 10838039001) overnight at 4°C. Plasmid transfections were performed using Lipofectamine 3000 (Thermo L3000015) according to the manufacturer’s protocol. The transfection amounts of different plasmids were optimized (Table S2). Plasmids generated in this study and their maps will be submitted to Addgene and also available on request. Cells were analyzed by microscopy imaging, Western Blot (WB) or other means 24 – 48 hr after plasmid transfection.

### Aggregate induction and inhibitor/chemical treatments

To examine protein aggregation under different conditions, cells were either untreated, mock treated or heat stressed at 42°C or treated with different chemicals for 12 – 16 hr. The proteasome inhibitor MG132 (Sigma M7449) was used at 10 μΜ final concentration. The autophagy inhibitor Bafilomycin A1 (Baf) (Sigma SML1661) was used at 100 nM. NaCl (Axil BIO-1111) was used at 200 mM for 4 hr to replicate the condition used for Sec body (SB) induction. The protein translation inhibitor cycloheximide (CHX) (Sigma C1988) was used at 20 μg/mL for 6 hr. 1,6-Hexanediol (Hex) (Sigma 88571) was used at 3.5% [v/v] and cells were imaged immediately by timelapse recording.

### HaloTag (Halo) and Immunofluorescence (IF) staining

Halo staining was performed according to the dye manufacturer’s protocol with minor modifications. Briefly, Janelia Fluor 549 (JF549) or JF646 ligand (Promega GA1110 or GA1120) was added to media to a final concentration of 15 – 50 nM and incubated for 15 – 30 min or indicated time. Then cells were washed once using fresh media (unless indicated otherwise), and either imaged directly or fixed using 4% [w/v] paraformaldehyde (PFA) (EMS 15710) diluted in phosphate buffered saline (PBS) (Thermo 14190250) for 20 min at room temperature (RT). For IF staining of proteins except from calreticulin (CRT), live cells were washed once using fresh media and fixed using 4% [w/v] PFA. After three PBS washes, cells were permeabilized in PBS + 0.1% [v/v] Triton X-100 (Thermo 85111) for 10 min and blocked in PBS + 0.1% [v/v] Tween 20 (Sigma P1379) (PBST) + 1% [w/v] BSA (Sigma A2153). Cells were incubated with primary antibodies overnight at 4°C in PBST + 1% [w/v] BSA. Then after four washes with PBS, cells were incubated with fluorophore-conjugated secondary antibodies in the same diluent for 1 hr at RT. Again, cells were washed four times using PBS before microscopy imaging. For IF staining of CRT, the following modifications were made: cells were instead fixed using 2% [w/v] PFA + 1.5% [w/v] glutaraldehyde (GA, Sigma 5882) at 37°C; after PBS washes to remove fixatives, cells were incubated for 10 min at RT with 0.1% [w/v] sodium borohydride (NaBH_4_, Sigma 452882) in PBS to reduce GA; primary antibody incubation was reduced to 2 hr at 4°C. Antibodies used in this study are listed in Table S2.

### RNA fluorescence *in situ* hybridization (FISH)

RNA FISH was performed according to a published protocol (Jain et al., 2016) with minor modifications. Briefly, cells were fixed and permeabilized as in IF, and then washed thrice in PBS and twice in SSC wash buffer (2x SSC buffer + 15% [v/v] formamide), in which the 2x SSC buffer (300 mM NaCl, 30 mM sodium citrate, pH = 7.0) was diluted from 20x SSC (Sigma S6639) using UltraPure DNase/RNase-Free Distilled Water (Thermo 10977015). Then Alexa Fluor 647 (AF647)-conjugated oligo(dT)_30_ purchased form IDT was diluted to a final concentration of 1 ng/μL in FISH buffer, which consists of SSC wash buffer + 10% [w/v] dextran sulfate (Sigma D8906) + 2 mM vanadyl-ribonucleoside complex (NEB S1402) + 0.02% [w/v] BSA, + 1 mg/mL tRNA (Sigma 10109541001), and used to stain cells at 37°C overnight. After incubation, cells were washed twice in SSC wash buffer and twice in PBS, re-fixed in 4% [w/v] PFA for 10 min at RT, and washed thrice in PBS before imaging.

### Microscopy imaging

Wide field images were taken on a EVOS M5000 microscope using a 40x air objective. Confocal images were taken on Yokogawa CSU-W1 spinning disk microscope (installed on Nikon Eclipse Ti-E inverted microscope body) with a CFI Apochromat λ 100x oil objective (NA = 1.4), Cargille Type 37 immersion oil (Cargille Labs 16237-16) and Photometrics Prime 95B Scientific CMOS cameras. Z-stacks were taken using a Piezo stage (Piano Z from Physik Instrumente) every 0.1 μm for a total thickness of 5 or 10 μm except for Video S9, in which the step size was 0.5 μm. To enable Structured Illumination Microscopy (SIM), the Live-SR super-resolution module (Roper Scientific) was engaged except for Figure 4, 5E, G, 6D, E, S6A, B, S8, S9C and E. For live cell imaging, the environment chamber was stabilized to 37°C and CO_2_ was provided by LCI FC-5N CO_2_ mixer (Live Cell Instrument) and calibrated to maintain normal pH in cell culture media. For imaging heat stressed cells, the on-stage incubator (Live Cell Instrument CU-501) was heated to and stabilized at 42°C. To avoid drifting, cells were pre-treated in a 42°C CO_2_ incubator for 15 min before they were transferred to the microscope and allowed 30 min to further stabilize before imaging starts.

### Correction of chromatic aberration, image analysis and statistics

Chromatic aberration of confocal images along the z direction was calibrated using 0.1 μm Tetraspec Microspheres (Thermo T7279) and corrected in ImageJ/Fiji (Schindelin et al., 2012) using a custom Jython script. 3D reconstruction and analysis of z-stacks were performed in Imaris whereas single z-slice images were analyzed using custom ImageJ Jython scripts. Measurements made in Imaris and ImageJ were analyzed using custom R scripts.

### Quantification of coaggregates

For quantification of the (volume) percentage of SEC16A inclusions with TDP43 aggregation, TDP43/SEC16A coaggs and SEC16A inclusions without TDP43 were respectively identified in Imaris using its Labkit machine learning plugin (Arzt et al., 2022). For other quantifications, inclusions of two different proteins, *e.g.* SEC16A and Hsp104^DWB^, were respectively identified in Imaris by manual thresholding based on local contrast, and the mutual overlap volumes between pairs of aggregates were measured. A SEC16A inclusion is categorized as in colocalization with Hsp104^DWB^ if the overlap volume is no less than 80% of the former’s volume.

### Fluorescence recovery after photobleaching (FRAP) assay

Aggregates were imaged for 11 frames before photobleaching by high power laser in part or in full. Then the aggregates were imaged continuously for at least 2.5 min and quantified in ImageJ using a custom Jython script.

### RNA inhibition (RNAi)

TriFECTa RNAi Kits including predesigned dicer-substrate small interference RNAs (DsiRNAs) were purchased from IDT. The sequences of DsiRNAs are listed in Table S2. Each DsiRNA was co-transfected at a final concentration of 10 nM with plasmids using Lipofectamine 3000 according to the manufacturer’s protocol. Briefly, DsiRNAs were immediately added after mixing Lipofectamine 3000 and P3000-bound plasmids.

### Western blot (WB)

Cells cultured in each 35 mm dish were dissolved in 200 – 300 μL RIPA buffer (Thermo 89900) + 2 mM PMSF (Sigma P7626) + 3 μL Protease Inhibitor Cocktail (PIC, Sigma P8340) with regular swirling for 10 min. The protein concentrations of cell lysates were measured using BCA assay (Thermo 23225) and equal amounts of proteins were loaded onto 4 – 15 % polyacrylamide gradient gels (Bio-Rad 4561084) and fractionated by SDS-polyacrylamide gel electrophoresis (SDS-PAGE). Then proteins were electroblotted onto nitrocellulose membranes (BIO-RAD 1704158). After blocking in Odyssey Blocking Buffer (PBS, LI-COR 927), membranes were incubated sequentially with primary and secondary antibodies in Odyssey Blocking Buffer mixed with an equal volume of PBS + 0.2% [v/v] Tween 20. After each incubation, membranes were washed four times in PBST. Tween 20 was removed by rinsing in PBS before detecting the fluorescence of secondary antibodies using a LI-COR Odyssey CLx Scanner. The fluorescence of protein bands was quantified in ImageStudioLite (Bio-Rad).

### Aggregate purification

HEK293T cells cultured in two T225 flasks (Thermo 159934) were respectively transfected with plasmids carrying Hsp104^DWB^-mEGFP-FLAG and, as a control, Hsp104^DWB^-mEGFP. 24 hours after transfection, both flasks were transferred to a 42°C CO_2_ incubator for 12 hr to exacerbate protein aggregation. Then cells were gently washed once in PBS and harvested by washing off using PBS. The pellets from centrifugation at 120 g for 5 min was washed again using PBS and frozen in liquid nitrogen for future use. When needed, the frozen pellets were thawed at 4°C, each lysed by incubation in 1000 μL RIPA buffer + 2 mM PMSF + 60 μL PIC + 112.5 Kunitz Unit/mL DNase I (RNase-free, NEB M0303) + 2.5 mM MgCl_2_ (Sigma 5985-OP) with occasional pipette aspiration, and a brief centrifugation at 150 g for 5 min was conducted to remove cell debris. A small aliquot of the supernatant was used for BCA assay of protein concentration whereas the rest was loaded onto sucrose gradient layers of 20% and 50% [w/v] sucrose, each dissolved in 1500 μL RIPA buffer. Centrifugation at 60,000 g for 30 min on an ultracentrifuge (Thermo Sorvall WX100) enriched Hsp104^DWB^-mEGFP(-FLAG)-labelled aggregates at the interface between 20 and 50% [w/v] sucrose. This layer was extracted through needle and syringe to yield ∼ 750 μL of sample, which in turn was mixed with 200 μL of pre-equilibrated anti-FLAG M2 resin (Sigma A2220), rotated at 4°C for 1 hr, and washed three times with RIPA buffer to immunoprecipitate Hsp104^DWB^-mEGFP-FLAG-labelled aggregates. In the end, aggregates were eluted in urea elution buffer (8M urea, 20 mM Tris pH=7.4, 100 mM NaCl). At each step, the presence of Hsp104^DWB^-mEGFP-FLAG-labelled aggregates was checked by microscopy, and aliquots were saved for analysis by SDS-PAGE and Coomassie Brilliant Blue staining or WB.

### Liquid chromatography - tandem mass spectrometry (LC – MS/MS)

Proteins were reduced by 10 mM TCEP for 20 min at RT and alkylated with 55 mM 2-chloroacetamide (CAA) in the dark for 30 min. Then, the proteins were digested with 2 μg endoproteinase LysC for 3 hr followed by 2 μg trypsin at 37°C overnight, and the digestion was terminated by adding trifluoroacetic acid (TFA) to a final concentration of 1% [v/v]. Subsequently, the peptides were desalted using C18 HLB cartridges (Waters), dried by centrifugal evaporation, and resuspended in 25 μl TEAB, pH = 8.5. The samples were differentially labelled with TMT10-plex isobaric tandem mass tags (Thermo) at 25°C overnight, and the reaction was quenched by addition of 30 μl of 1 M ammonium formate, pH = 10 before pooling all samples into a low-binding microfuge tube. The pooled samples were desalted and fractionated on a self-packed spin column filled with C18 beads (Dr Maisch) in the step gradient of 14%, 24% and 60% [v/v] acetonitrile dissolved in 10 mM ammonium formate, pH=10. Each fraction was dried by centrifugal evaporation and further washed and dried twice by addition of 60% [v/v] acetonitrile + 0.1% [v/v] formic acid to remove residual ammonium formate. Then the samples were resuspended in 10 μl of 2% [v/v] acetonitrile + 0.06% [v/v] TFA + 0.5% [v/v] acetic acid and transferred to an autosampler plate. Online chromatography was performed using an EASY-nLC 1000 liquid chromatography system (Thermo) with a single-column setup and 0.1% [v/v] formic acid and 0.1% [v/v] formic acid + 99% [v/v] acetonitrile as mobile phases. The fractionated samples were injected and separated on an Easy-Spray reversed-phase C18 analytical column (Thermo) with 75 μm inner diameter, 50 cm length and 2 μm particle size. The column was maintained at 50°C in 2 - 33% [v/v] acetonitrile gradient for over 55 min, followed by an increase to 45% [v/v] over the next 5 min, and then to 95% [v/v] over 5 min. The final mixture was maintained on the column for 4 min to elute all remaining peptides. The total run duration for each sample was 70 min at a constant flow rate of 300 nL/min. MS data were acquired on a Q Exactive HFX mass spectrometer (Thermo) using data-dependent mode. Samples were ionized under 2.1 kV and 300°C at the nanospray source. Positively charged precursor signals (*i.e.* MS1) were detected using an Orbitrap analyzer set to a resolution of 60,000, automatic gain control (AGC) target of 3,000,000 ions, and max injection time (IT) of 50 ms. Precursors with charges 2 - 7 of the highest ion counts in each MS1 scan were further fragmented using higher-energy collision dissociation (HCD) at 36% normalized collision energy. The fragment signals (*i.e.* MS2) were analysed by the Orbitrap analyzer at a resolution of 7,500, AGC of 100,000 and max IT of 100 ms. Data-dependent (MS2) scans were acquired in Top40 mode. To avoid re-sampling of high abundance peptides, the precursors used for MS2 scans were excluded for 30 sec.

### Proteomics data analysis

Proteins were identified using Proteome Discoverer v2.3 (Thermo). Briefly, the raw mass spectra were searched against human primary protein sequences retrieved from Swiss-Prot (June 11 2019 release) (Boutet et al., 2007) with carbamidomethylation on Cys and TMT10-plex on Lys and N-terminus set as fixed modifications, and deamidation of Asn and Gln, acetylation of N-terminus, and Met oxidation as dynamic modifications. Trypsin/P was set as the digestion enzyme and allowed up to three missed cleavage sites. Precursors and fragments of mass errors within 10 ppm and 0.8 Da, respectively, were accepted, while reporter ions were at least 1 Da apart from each other. Peptides were matched to the spectra at a false discovery rate (FDR) of 1% (strict) and 5% (relaxed) against the decoy database and quantitated using TMT10-plex method. The search results were further processed using a custom R script, in which Gene Ontology (GO) enrichment was performed using clusterProfiler (Consortium, 2015, Yu et al., 2012). The SG/nucleolus proteome, IDR-containing proteins and LLPS substrates were retrieved from previous publications and databases (Marmor-Kollet et al., 2020, Thul et al., 2017, Piovesan et al., 2021, Wang et al., 2022).

### Retention Using Selective Hooks (RUSH) assay for measuring ER-to-Golgi transport

RUSH was performed as previously described with minor modifications (Figure S7A) (Boncompain et al., 2012). Briefly, the RPE-1 cell culture media was substituted with custom made biotin-free DMEM:F12 media (Cell Culture Technologies) + 10% [v/v] FBS from seeding (1 d before transfection). Then a plasmid carrying both the ER anchor ss-Strep-KDEL and the reporter ManII (aa1 – 116 only)-SBP-mCherry or TNFα-SBP-mCherry was co-transfected with TDP43 and SEC16A constructs using Lipofectamine 3000. To release cargo retention, D-biotin (Thermo B20656) was added to a final concentration of 40 μM. For tracking ERES in timelapse imaging, SEC16A inclusions were identified in ImageJ using ilastik machine learning plugin (Berg et al., 2019) and then tracked by Trackmate (Ershov et al., 2022). The trajectories were analyzed by custom R scripts. Only SEC16A inclusions that enriched transport cargo (RUSH-TNFα) and tracked for no less than 10 frames (∼ 15 sec) were used for cross-correlation (CC) analysis.

### AP21967-induced TDP43-ERES coaggregation

3xFKBP-TDP43-mNG and 3xFRB*-Halo-SEC16A were co-transfected with the RUSH-TNFα construct. 1 μM rapamycin analog AP21967 (AP, Takara 635056) was added ∼ 18 hr post-transfection to treat the cells for 6 – 10 hr to induce coaggregation.

### Differentiation of motor neurons (MNs) from stem cells

The human embryonic stem cells (hESCs) and induced pluripotent stem cells (hiPSCs) used in this study are: NDS00268, NDS00269 and NDS00270 (ALS patients) and BJ, GM23720 and H9 (non-ALS). NDS00268, NDS00269, NDS00270 and GM23720 were obtained from the NIGMS Human Genetic Cell Repository at Coriell Institute, whereas H9 was purchased from Corning. The induction of MNs from stem cells were performed according to previous publications (Hor et al., 2018, Hor et al., 2021) and illustrated in Figure S7A. In brief, stem cells were cultured on Matrigel (Corning 354230)-coated plates in StemMACS iPS-Brew XF (Miltenyi Biotec 130-104-368). The cell culture media was changed every other day and cells were passaged at a split ratio between 1: 6 to 1: 10 every 6 - 8 days. On the day of differentiation (D0), stem cells were digested with Accutase (Nacalai Tesque 12679-54) for 5 - 10 min to acquire single-cell suspensions. Accutase was removed by centrifugation and PBS wash. Then cells were seeded at 70% confluency on dishes pre-coated with Matrigel and cultured in StemMACS iPS-Brew XF + 5 μM Y-27632 (STEMCELL Technologies 72304). On D1, the media was replaced with Neural Induction Media (NIM) comprising the Neural Media (NM) + 0.5 μM LDN193189 (STEMCELL Technologies 72147) + 4.25 μM CHIR99021 (Miltenyi Biotec 130106539), where NM is a mixture of 50% [v/v] DMEM:F12 + 50% [v/v] MACS Neuro Media (Miltenyi Biotec 130-093-570) + 0.5x GlutaMax (Thermo 35050061) + 1x NEAA (Thermo 11140050) + 1x N-2 (Thermo 17502048) + 1x MACS NeuroBrew-21 (Miltenyi Biotec 130-093-566). On D3, 1 μM retinoic acid (RA, Sigma R2625) was supplemented to the media, which was then changed every 2 - 3 days until D7 or when cells became confluent. At this point, stem cells have differentiated into MN progenitors, which can be passaged at a ratio between 1: 2 and 1: 3 onto new Matrigel-coated dishes. These progenitor cells were cultured in NIM + 1 μM RA + 5 μM Y-27632 and the media was changed every day. From D11, MN differentiation media, *i.e.* NM + 1 μM RA + 1 μΜ Purmorphamine (Miltenyi Biotec 130-104-465), was added and changed every other day until D17. Then MN maturation media, *i.e.* NM + 10 ng/mL GDNF (Miltenyi Biotec 130-129-547) + 10 ng/mL BDNF (Miltenyi Biotec 130-093-811) + 200 μM L-ascorbic acid (Sigma A5960), was added and changed every other day until MNs were ready on D28. The quality of MNs was checked by morphology and IF staining of (motor) neuron markers including NeuN/RBFOX3, NFM and ChAT.

### Immunofluorescence staining of post-mortem ALS patient samples

The post- mortem samples of sALS patients were acquired from the Johns Hopkins/Temple University ALS Postmortem Core in form of formalin-fixed paraffin-embedded (FFPE) tissue sections of approximately 20 mm x 10 mm x 5 μm in dimensions. The samples were deparaffinized by incubating thrice in fresh Xylene (Sigma 534056) and rehydrated by incubating twice in ethanol and once in 90% and 70% [v/v] ethanol. Each incubation time is 5 min. Then the samples were boiled in epitope retrieval solution (IHC World IW-1100) in an autoclave (Hirayama HV-50) using the liquid cycle (121°C 15 min), washed twice with PBS, and permeabilized for 10 min in PBS + 0.4% [v/v] Triton X-100. After three PBS washes, the samples were blocked in serum free protein block (Agilent X090930-2) for 1 hr at RT, and incubated with primary antibodies diluted in background reducing antibody diluent (Agilent S302283-2) overnight at RT. After three PBS washes, the samples were incubated with diluted secondary antibodies for 2 hr at RT and washed with PBS. The nuclei were stained by incubation with 1 μg/mL DAPI (Sigma MBD0015) in PBS for 30 min at RT followed by three more PBS washes. The stained sections were preserved in Prolong Gold antifade mountant (Thermo P36930) with coverslips sealed by transparent nail polish (Beauty Language). For image quantification, only cells positive for anti-NeuN immunostaining were counted.

## Author Contributions

H.W. and R.L. designed the study and wrote the manuscript. H.W. conducted and performed data analysis for most experiments. L.C.W. performed LC-MS/MS. B.M.S. determined the solid/liquid-like states of aggregates. D.L. derived MNs from stem cells. J.Z. designed RNAs for targeting genes of interest. K.M.G. and K.W. prepared post-mortem tissue sections from ALS patient samples and helped optimize immunofluorescence staining protocol. R.G. assisted with molecular cloning and cell staining. C.J.J.Y., R.M.S. and L.W.O. contributed to experiment designs. R.L. supervised the project.

## Acknowledgements

The authors would like to thank J. Shorter (UPenn) for recommending the DWB mutations for stable Hsp104-aggregate binding and generously sharing a Hsp104^DWB^construct. The authors thank P. Kanchanawong and K. Paramasivam for kindly sharing FP constructs. The authors thank H.T. Ong and J.F.L. Chin from MBI Microscopy core, Singapore Microscopy and Bioimage Analysis (SIMBA), for help in correction of chromatic aberration in confocal imaging, and MBI Wet Lab Core for providing essential lab equipment. The authors are grateful to G. Thibault (NTU), B. Burke (A*STAR), T. Hiraiwa, G. Jedd (TLL), P.T. Matsudaira, J. Hu, M. Yao, W. Huang, L. Ruan (JHU), H. Wang, R. Das, T. Chew, Y. Tee, M. Pan, S. Ramachandran (JHU), B. S. Wong, K. Seah and Y. Lee for constructive discussions. This work was supported by R. Li’s startup grant (A-0007081-00-00) and a Tier 3 grant (A-0006324-03-00) received from Singapore Ministry of Education, and by the A*STAR core funding and Singapore National Research Foundation (under its NRF-SIS “SingMass” scheme) endowed to R.M. Sobota and L.C. Wang.

## Notes

MS proteomics data have been deposited to the Japan ProteOme STandard Repository (jPOSTrepo) under identifier 9443.

## Abbreviations

3D: 3 dimension
aa: Amino acid
AF: Alexa Fluor
AGC: Automatic gain control
ALS: Amyotrophic Lateral Sclerosis
Baf: Bafilomycin A1
BCA: Bicinchoninic acid
BDNF: Brain Derived Neurotrophic Factor
C9: C9ORF72
CAA: 2-chloroacetamide
ChAT: Choline acetyltransferase
CHX: Cycloheximide
Coagg: Coaggregate
COPII: Coat Protein Complex II
CRT: Calreticulin
DMEM: Dulbecco’s Modified Eagle Medium
DNA: Deoxyribonucleic acid
DsiRNA: Dicer-substrate siRNA
DWB: Double Walker B
EGFP: Enhanced green fluorescent protein
ER: Endoplasmic reticulum
ERES: ER-exit site
FACS: Fluorescence-activated cell sorting
FBS: Fetal bovine serum
FDR: False discovery rate
FISH: Fluorescence in situ hybridization
FP: Fluorescent protein
FRAP: Fluorescence recovery after photobleaching
FUS: Fused in sarcoma
GDNF: Glial cell line-derived neurotrophic factor
GM130: Golgi matrix protein 130 kDa
Halo: HaloTag
HCD: Higher-energy collision dissociation
HEK293T: Human embryonic kidney 293T
hESC: Human embryonic stem cell
Hex: 1,6-Hexanediol
hiPSC: Human induced pluripotent stem cell
hnRNP A1: Heterogeneous nuclear ribonucleoprotein A1
Hsp: Heat-shock protein
Htt: Huntingtin
IDR: Intrinsically disordered region
IF: Immunofluorescence
iMN: Induced MN
INS-1: (Rat) Insulinoma-1
IP: Immunoprecipitation
IPOD: Insoluble protein deposit
IT: Injection time
JF: Janelia Fluor
JUNQ: Juxta-nuclear quality control compartment
mEGFP: Monomeric EGFP
mNG: Monomeric NeonGreen
NeuN: Neuronal nuclei antigen
G3BP1: Ras GTPase-activating protein-binding protein 1
LC: Liquid chromatography
LLPS: Liquid-liquid phase separation
ManII: Mannosidase II
MN: Motor neuron
MS: Mass-spectrometry
MTOC: Microtubule organization center
NBD: Nucleotide-binding domain
NEAA: Non-essential amino acids
NFM: Neurofilament medium polypeptide protein
NM: Neural media
NIM: Neural Induction Media
NLS: Nuclear localization signal
NTD: N-terminal domain
PBS: Phosphate buffered saline
PBST: PBS + 0.1% [v/v] Tween 20
PCR: Polymerase chain reaction
PD: Parkinson’s Disease
PFA: Paraformaldehyde
PIC: Protease inhibitor cocktail
PMSF: Phenylmethylsulfonyl fluoride
PolyA: Poly-adenine
PS-1: Presenilin-1
RA: Retinoic acid
RBFOX3: RNA binding protein fox-1 homolog 3
RIPA: Radioimmunoprecipitation assay
RNA: Ribonucleic acid
RNAi: RNA interference
RPE-1: Retinal pigment epithelial-1
RRM: RNA-recognition motif
RT: Room temperature
RUSH: Retention Using Selective Hooks
sALS: sporadic ALS
SB: Sec body
SBP: Streptavidin-binding protein
SDS: Sodium dodecyl sulfate
SDS-PAGE: SDS-polyacrylamide gel electrophoresis
SG: Stress granule
SHH: Sonic Hedgehog
SIM: Structured Illumination Microscopy
siRNA: Small interfering RNA
SOD1: Superoxide dismutase 1
SR: Super-resolution
ss: (Secretory) signal sequence
SSC: Saline-sodium citrate
TCEP: tris(2-carboxyethyl)phosphine
TDP43: TAR DNA-binding protein 43 kDa
TEAB: Triethylammonium bicarbonate
TFA: Trifluoroacetic acid
TNFα: Tissue necrosis factor α
tRNA: transfer RNA
VC: Vitamin C (L-ascorbic acid)
WB: Western blot
WT: Wild-type

---

**Figure S1.**
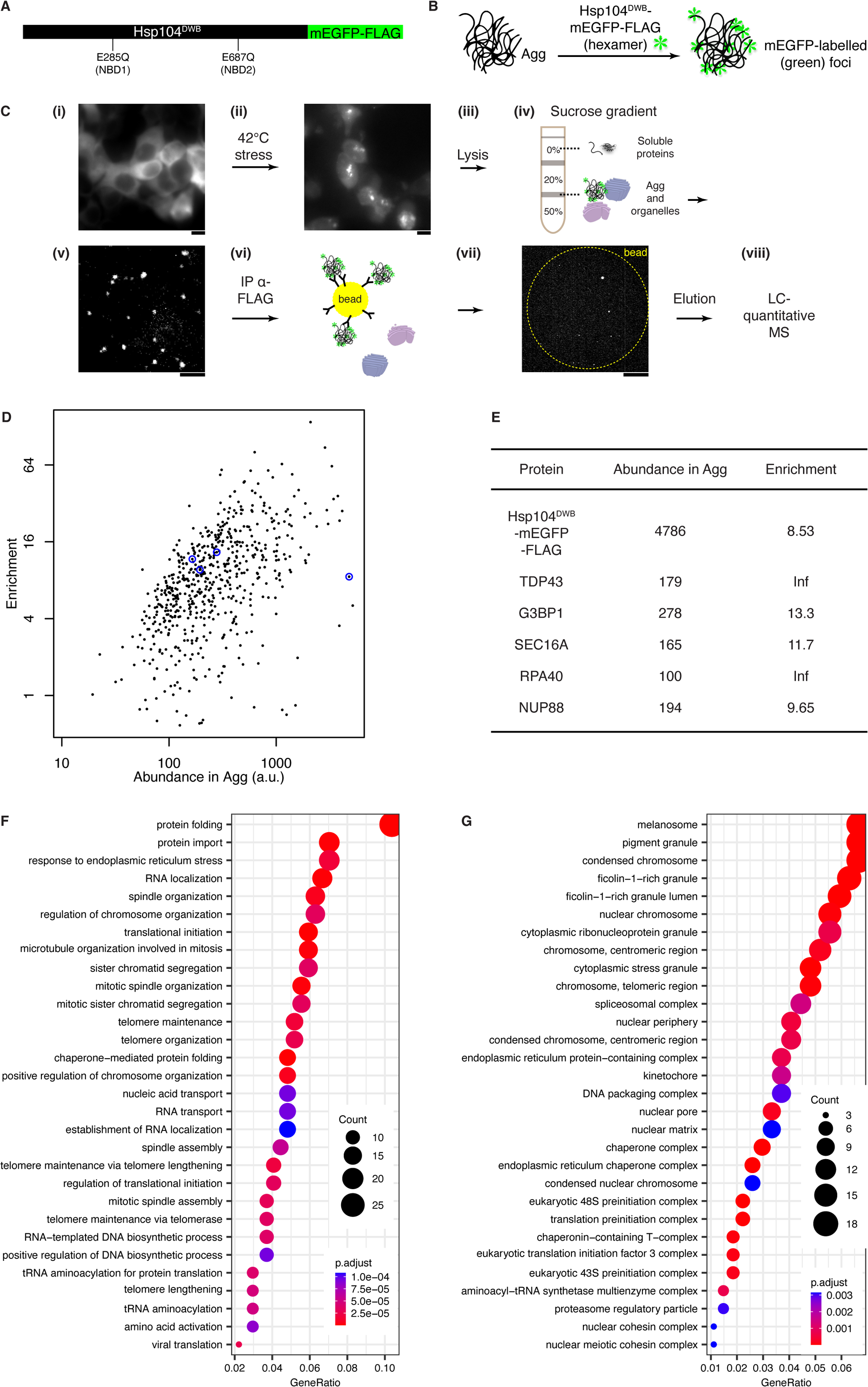
Hsp104^DWB^ enables affinity-based aggregate purification. (**A**) Schematic diagram of Hsp104^DWB^-mEGFP-FLAG showing the locations of double Walker B (DWB) mutations in both nucleotide binding domains (NBDs). (**B**) Working scheme of aggregate labelling. Multiple Hsp104^DWB^-mEGFP hexamers bind misfolded proteins in aggregates but not release to form bright foci. (**C**) Workflow of protein aggregate purification by sucrose gradient centrifugation and immunoprecipitation. (i) HEK293T cells transfected with Hsp104^DWB^-mEGFP-FLAG were (ii) heat stressed at 42°C for 12 hr and (iii) lysed by RIPA buffer. (iv and v) The lysate was fractionated by sucrose gradient centrifugation to enrich protein aggregates at the interface between 20% and 50% [w/v] sucrose layers. (vi and vii) Then Hsp104^DWB^-bound aggregates were immunoprecipitated (IP) using anti-FLAG beads and (viii) sent for liquid chromatography (LC) - quantitative mass-spectrometry (MS). Aggregates were imaged by Hsp104^DWB^-mEGFP-FLAG fluorescence at the various steps. The images in (i) and (ii) were captured by a widefield epifluorescence microscope whereas (v) and (vii) are the max intensity projections of confocal z-stacks. Each scale bar represents 10 μm. The yellow dashed circle in (vii) demarcates an anti-FLAG bead in the transmission light channel. (**D**) Scatter plot showing the abundance of MS-identified proteins in purified aggregates (plotted on a log_10_ scale) versus fold enrichment in aggregates (*i.e.* ratio between aggregate and IP control, plotted on a log_2_ scale). The positions of Hsp104^DWB^-mEGFP-FLAG and proteins later investigated are circled with values tabulated in (E). a.u.: arbitrary unit. N = 1. (**E**) Table showing the abundance and fold enrichment in aggregate of several proteins investigated in this study. Infinity (Inf) arose as some proteins were detected only in aggregates but not IP control. (**F and G**) GO analysis of the biological processes (F) and subcellular locations (G) of proteins enriched in aggregates. GeneRatio: fraction of aggregate proteins annotated in the respective GO term; Count: number of aggregate proteins annotated in the respective term; p.adjust: adjusted p-value of hypergeometric test.

**Figure S2.**
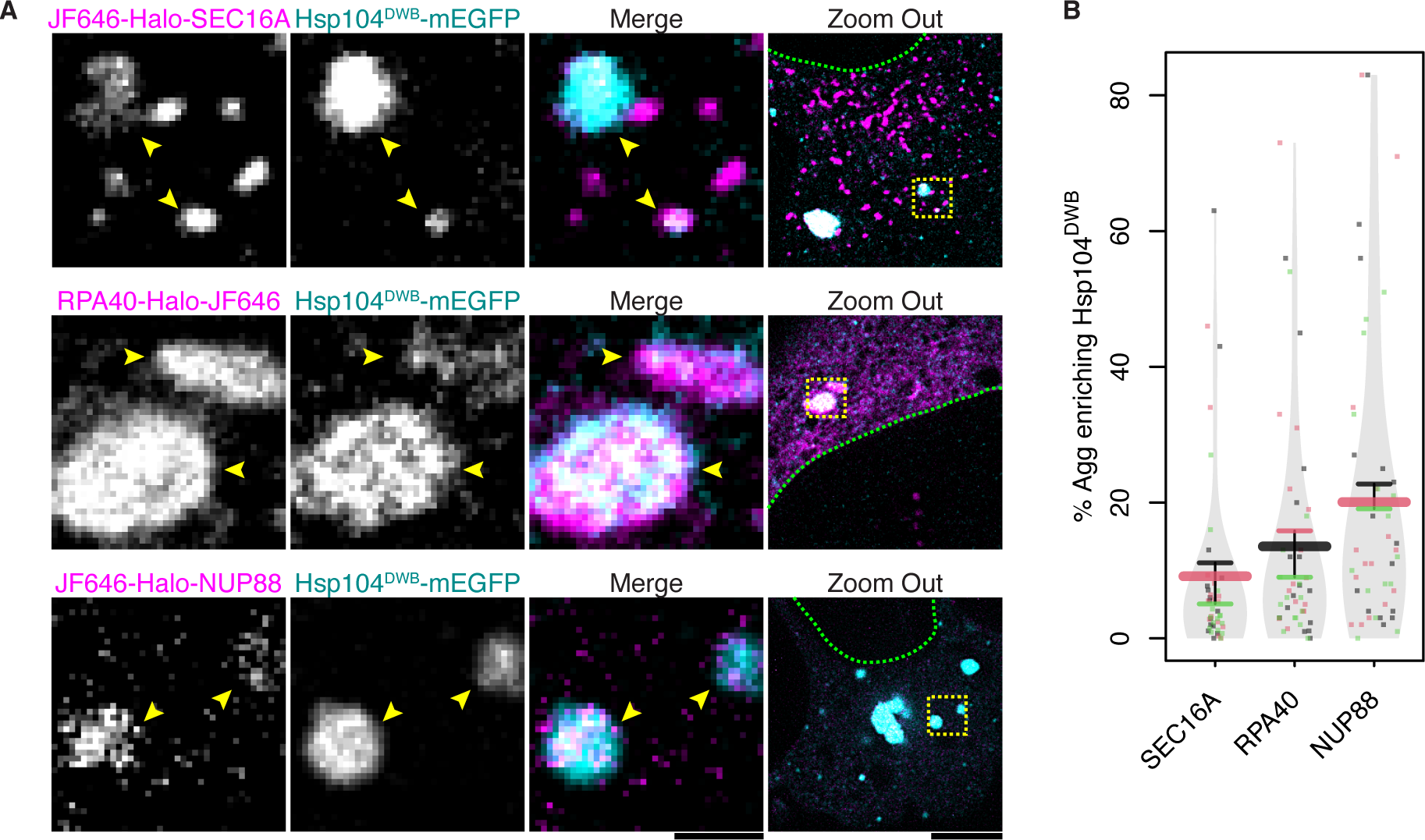
SEC16A, RPA40 and NUP88 form aggregates labeled with HSP104^DWB^-EGFP in HEK293T. (**A**) Representative images of aggregates formed by RPA40, SEC16A and NUP88 and their colocalization with Hsp104^DWB^. Similar to Figure 1G and H but in HEK293T cells stressed at 42°C for 12 - 16 hr. Arrowheads: Hsp104^DWB^-labelled aggregates; yellow boxes in “Zoom Out”: regions displayed in zoom-in views; green dashes: the nuclei. Scale bars except in “Zoom Out” represent 1 μm, and represent 5 μm in “Zoom Out”. (**B**) Quantification of the experiments as in (A) showing the percentage of RPA40, SEC16A and NUP88 aggregates labelled by Hsp104^DWB^ in each cell. N ≥ 3 x 15.

**Figure S3.**
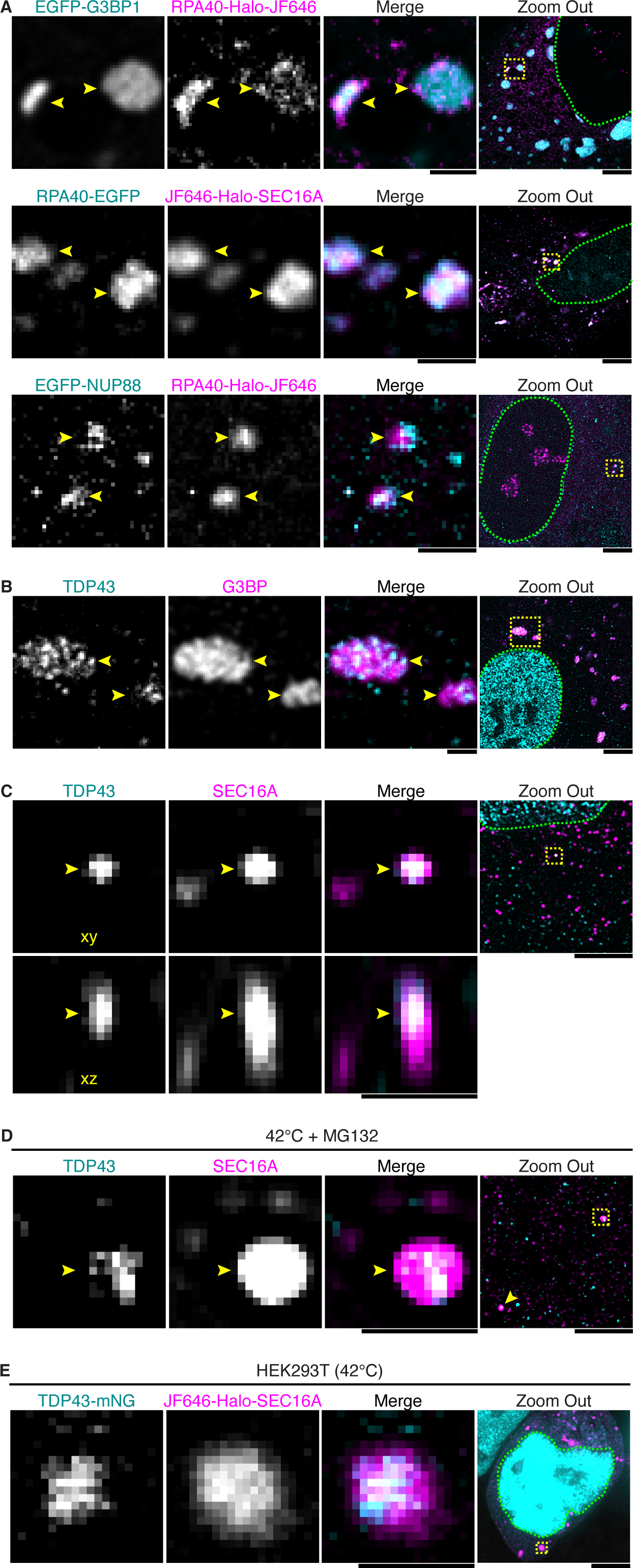
Proteins coaggregate with selective partners. (**A**) Representative images of the colocalization between RPA40 aggregates and aggregates formed by G3BP1, SEC16A or NUP88. Arrowheads: coaggregates; yellow boxes in “Zoom Out”: regions displayed in zoom-in views; green dashes: the nuclei. The quantification is shown in Figure 2H. Scale bars except in “Zoom Out” represent 1 μm, and represent 5 μm in “Zoom Out”. (**B**) Representative images of the immunofluorescence staining of TDP43 and G3BP (G3BP1/2) in cells forming SGs after incubation at 42°C for 16 hr. (**C**) Representative images of the immunofluorescence staining of TDP43 and SEC16A in cells forming coaggs after incubation at 42°C for 12 - 16 hr. (**D**) Representative images of the immunofluorescence staining of TDP43 and SEC16A in cells treated with 10 μM proteasome inhibitor MG132 and incubated for 12 - 16 hr at 42°C. (**E**) Representative images of TDP43/SEC16A coaggs formed in HEK293T cells after incubation for 12 - 16 hr at 42°C.

**Figure S4.**
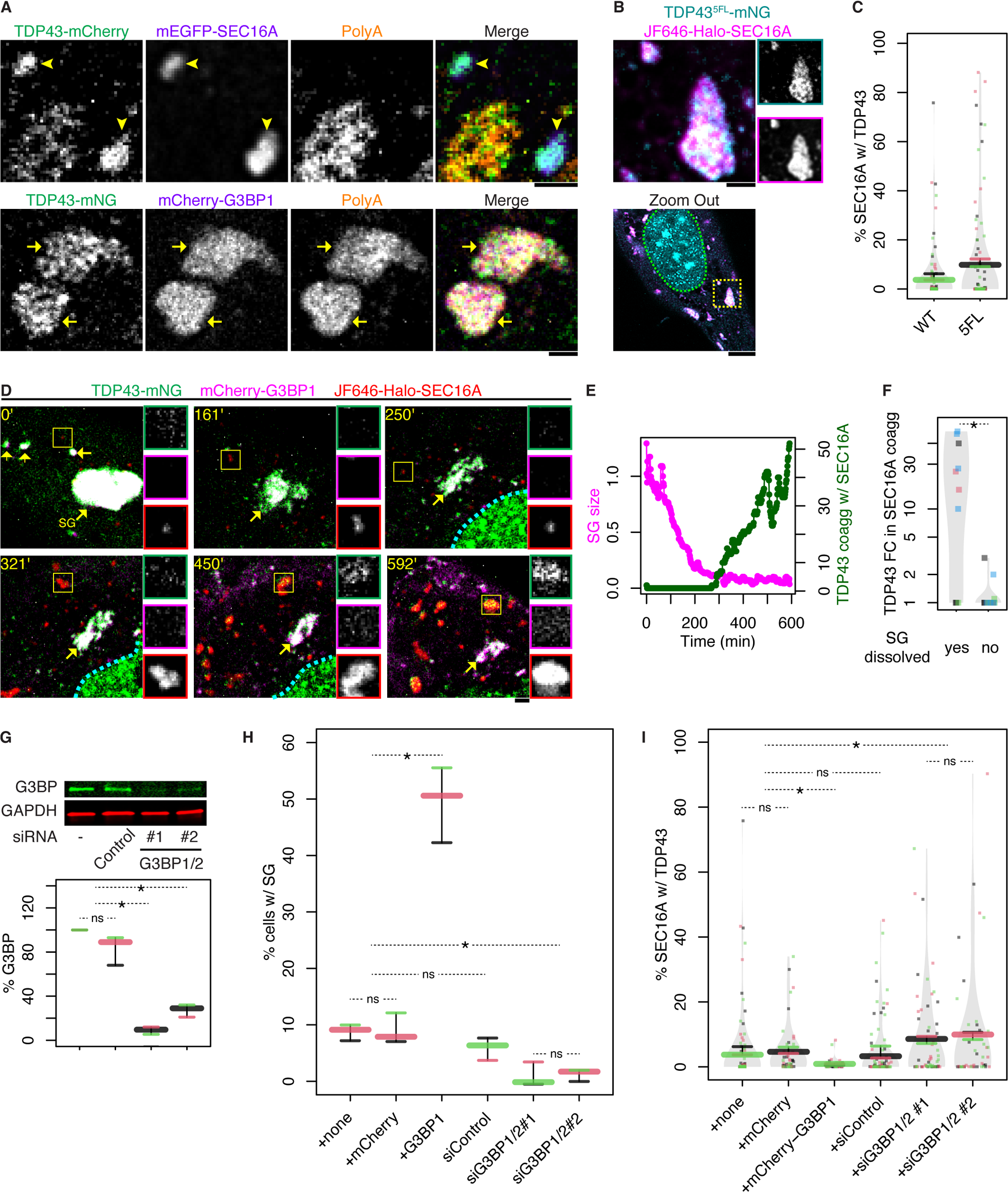
TDP43 partitions dynamically between SGs and coaggregates with SEC16A. (**A**) Representative images showing fluorescence in-situ hybridization (FISH) staining of polyA RNAs in cells containing TDP43/SEC16A coaggs (indicated by arrowheads) or SGs (indicated by arrows). PolyA RNAs were probed by oligo(dT)_30_ conjugated to AF647. Each scale bar represents 1 μm. (**B**) Representative images showing SEC16A coaggregation with TDP43^5FL^. In “Zoom out”, the yellow box marks the region shown in zoom-in views, and green dashes demarcate the nucleus. (**C**) Quantification of the volume percentage of SEC16A inclusions with wild-type (WT) TDP43 or TDP43^5FL^ aggregation in cells that expressed either construct. N ≥ 3 x 15. (**D**) Selected frames from a representative timelapse recording of SGs (indicated by arrows) and SEC16A inclusions in a cell shifted to 42°C for 60 min before imaging started. Insets show individual channels of the yellow boxed regions (SEC16A inclusion). (**E**) Quantification of the experiment in (D) showing the relative SG size and TDP43 intensity in coaggs with SEC16A over time (normalized to their respective initial values). (**F**) Quantification of the experiments as in (D) showing the fold changes (FCs) of TDP43 in SEC16A inclusions at the end of tracking of SG-containing cells, grouped by whether SGs were dissolved to below 10% of the initial size. 4 experiments were performed, and in each experiment ∼ 5 cells were tracked. The FC of each cell was plotted as a datapoint on log_10_ scale. (**G**) Western blot of G3BP1/2 in cells non-transfected or transfected with non-targeting (control) siRNA or different combinations of siRNAs against G3BP1 and G3BP2. Upper: representative blot; lower: quantification (N = 3). (**H and I**) Quantification of the percentage of cells forming SGs (H) and the volume percentage of SEC16A inclusions with TDP43 aggregation (I) after incubation at 42°C for 12 - 16 hr. Cells were either transfected with TDP43-mNG and Halo-SEC16A only or additionally with mCherry, mCherry-G3BP1, negative control siRNA, or different combinations of siRNAs against G3BP1 and G3BP2. SGs were stained by using FISH against polyA RNAs as in (A). N ≥ 3 x 15.

**Figure S5.**
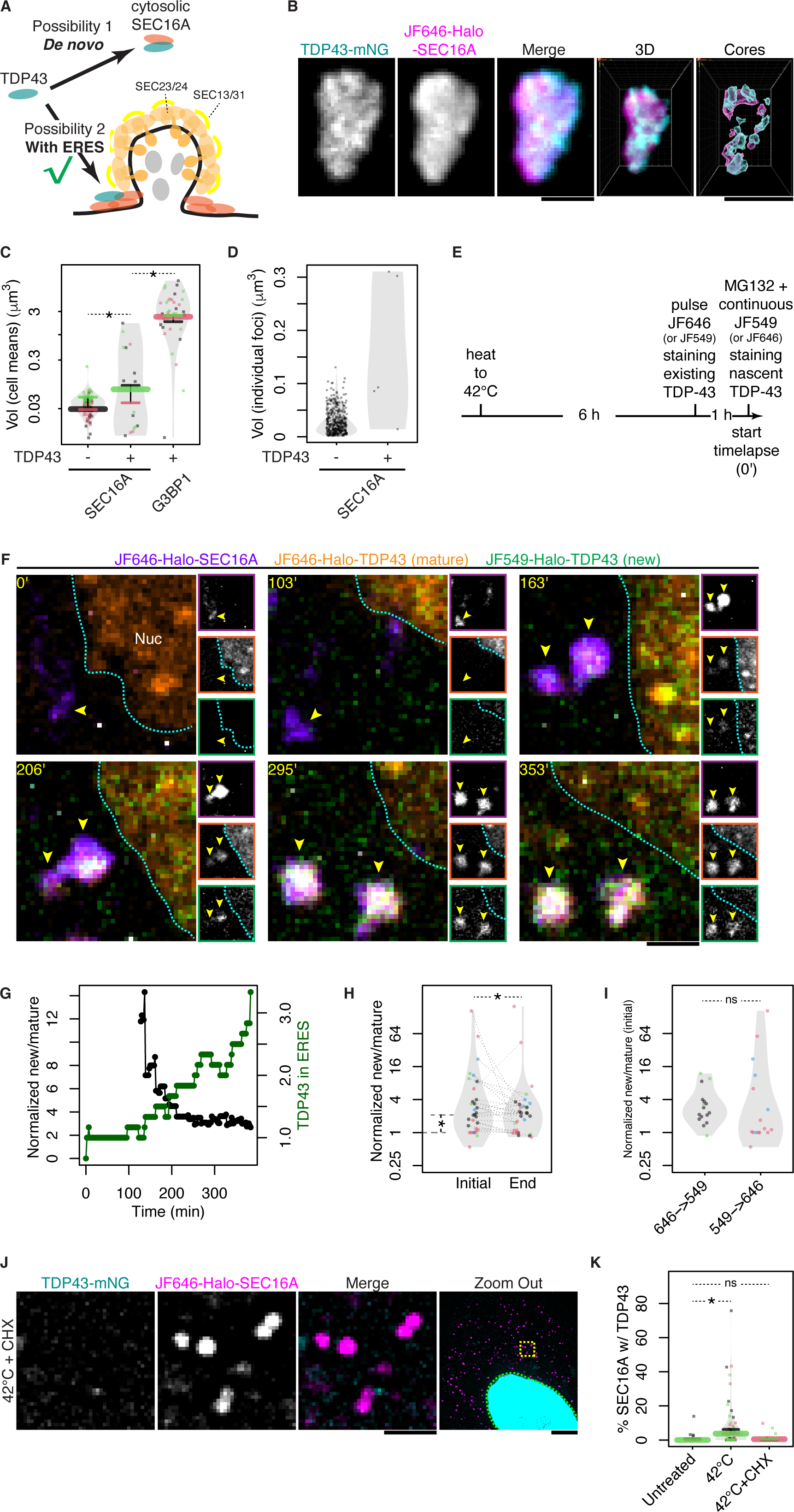
TDP43/SEC16A coaggs are ERES enriching nascent TDP43. (**A**) Schematic diagram showing two possible ways of TDP43/SEC16A coagg formation: *de novo* from cytosolic TDP43 and SEC16A, and TDP43 coaggregation with ERES. (**B**) Representative images of clusters of TDP43/SEC16A coaggs formed after 12 - 16 hr of incubation at 42°C. The 3D rendering shows a TDP43/SEC16A cluster viewed from top and “Cores” view annotates high intensity cores of TDP43 or SEC16A within the cluster. Each scale bar represents 1 μm unless otherwise indicated. (**C**) Quantification of the experiments as in Figure 2A and B comparing the volumes of SEC16A inclusions without TDP43, TDP43/SEC16A coaggs and TDP43-containing SGs. N ≥ 3 x 15. Volumes are plotted on a log_10_ scale. (**D**) Quantification of a cell with mild TDP43/SEC16A coaggregation (Figure 2B) showing the volumes of individual SEC16A inclusions with or without TDP43. Each data point represents a SEC16A inclusion. (**E**) Workflow of the pulse-chase of Halo-TDP43. Cells pre-stressed at 42°C for 6 hr were pulse-labelled for 30 min with the first dye, JF646 (or JF549), to visualize pre-existing TDP43 followed by dye wash off. Then 1 μM MG132 was used to inhibit TDP43 degradation followed by addition of the second dye, JF549 (or JF646), to continuously label newly synthesized TDP43. The second dye was kept in media. (**F**) Selected frames from a representative Halo-TDP43 pulse-chase. Pre-existing (mature) Halo-TDP43 was pulse-labelled by JF646 (orange) whereas newly synthesized (new) Halo-TDP43 was labelled continuously with JF549 (green). mEGFP-SEC16A (purple) was used to mark ERES. Insets show individual channels. Cyan dashes demarcate the nucleus (Nuc). (**G**) Quantification of the experiment in (F) showing the ratio between new and mature TDP43 that aggregated in ERES, normalized to the new/mature ratio of the whole cell, over time (black curve). To indicate the time of TDP43 aggregation in ERES, the relative intensity of TDP43 in ERES (normalized to its initial value) was shown (green curve). (**H**) Quantification of the experiments as in (F) showing the normalized new/mature TDP43 ratios upon aggregate formation (initial) and at the end of cell tracking (end), plotted on a log_2_ scale. 4 experiments were performed, and in each experiment ∼ 5 cells displayed TDP43 aggregation in ERES. The initial and end ratios of each cell are plotted as data points and, if the initial ratio exceeded 1, linked by gray segments. The asterisk between initial ratios and 1 indicates p ≤ 0.05 by one sample t-test, and the asterisk between linked initial and end ratios indicates p ≤ 0.05 by paired t-test. (**I**) Quantification of the experiments as in (F) showing the normalized new/mature TDP43 ratios upon aggregate formation, and grouped by the order of dye addition. 2 experiments were performed for each group, and in each experiment ∼ 5 cells displayed TDP43 aggregation in ERES. (**J**) Representative images of cells incubated at 42°C for 16 hr with cycloheximide (CHX) addition in the last 6 hr. In “Zoom Out”, the yellow box marks the region displayed in zoom-in views, green dashes demarcate the nucleus, and the scale bar represents 5 μm. (**K**) Quantification of the experiments as in (J), Figure 2B and 3A comparing the volume percentage of SEC16A inclusions with TDP43 aggregation in untreated cells or cells incubated at 42°C for 16 hr, without or with CHX treatment. N ≥ 3 x 15.

**Figure S6.**
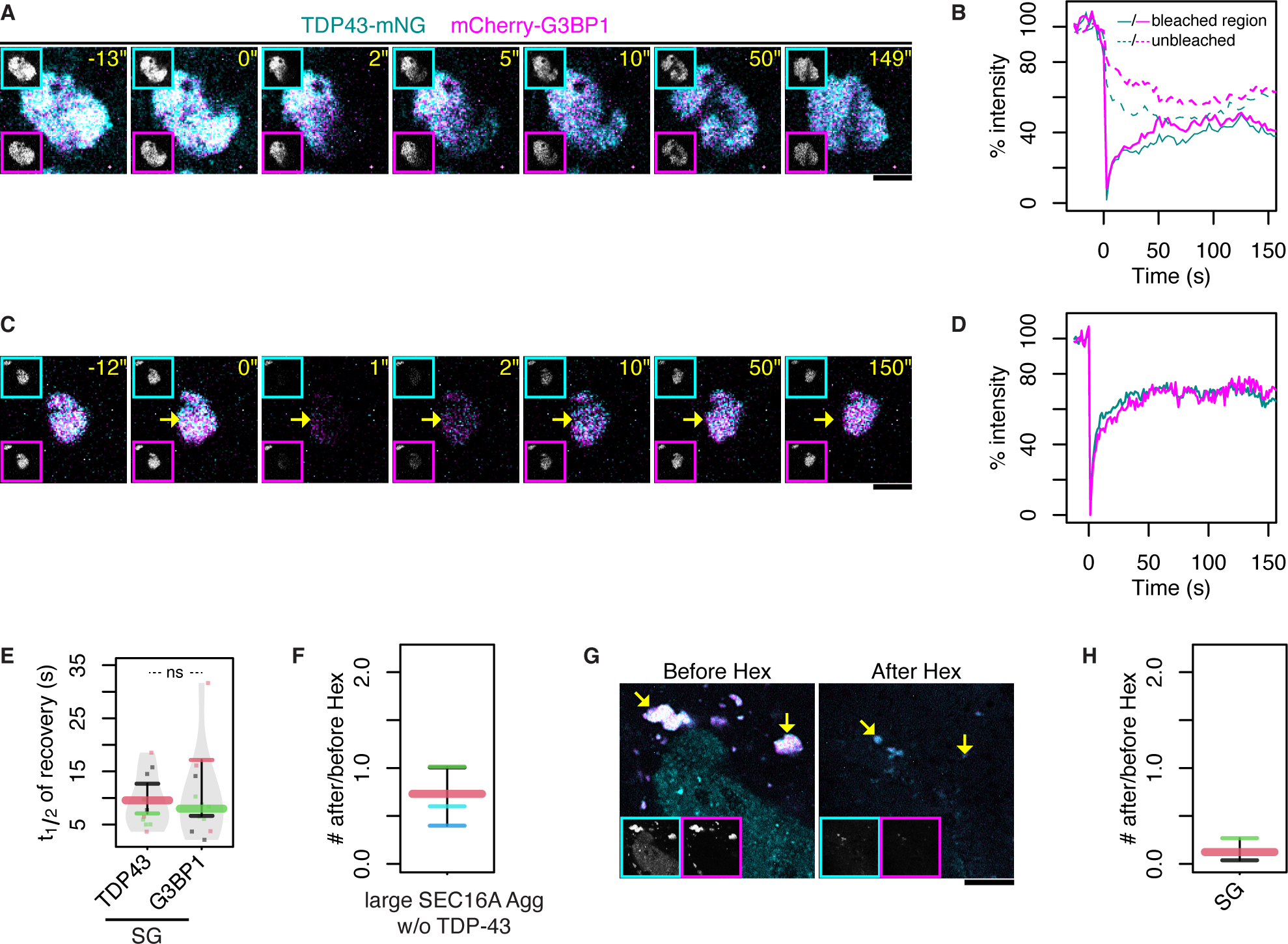
SGs and SEC16A inclusions without TDP43 are liquid-like. (**A**) Selected frames from a representative timelapse recording of a TDP43/G3BP1 coagg (TDP43-containing SG) after partial photobleaching. Insets show individual channels. (**B**) Quantification of the experiment in (A) showing the relative intensities of TDP43 (cyan lines) and G3BP1 (magenta lines) in bleached (solid lines) and unbleached regions (dashed lines) over time, normalized to their respective initial intensities. (**C**) Selected frames from a representative timelapse recording of a TDP43/G3BP1 coagg (SG) after photobleaching in its entirety. The position of the SG immediately before photobleaching is indicated by an arrow. (**D**) Quantification of the experiment in (C) showing the relative intensities of TDP43 (cyan line) and G3BP1 (magenta line) over time, normalized to their respective initial intensities. (**E**) Quantification of full FRAP experiments as in (C) showing the half time (t_1/2_) of fluorescence recovery for TDP43 and G3BP1 in SGs. 3 experiments were performed, and the average of ∼ 3 - 11 SG in each experiment was shown as a datapoint. (**F**) Quantification of the experiments in Figure 4G showing the after/before ratios of large ERES (of > 300 nm in radius). N ≥ 3 x 10. (**G**) Representative images showing SGs (indicated by arrows) in a cell before and after 3.5% [v/v] Hex treatment for 15 min. Insets show individual channels. The scale bar represents 5 μm. (**H**) Quantification of the experiments as in (G) showing the after/before ratios of SGs. N ≥ 3 x 10.

**Figure S7.**
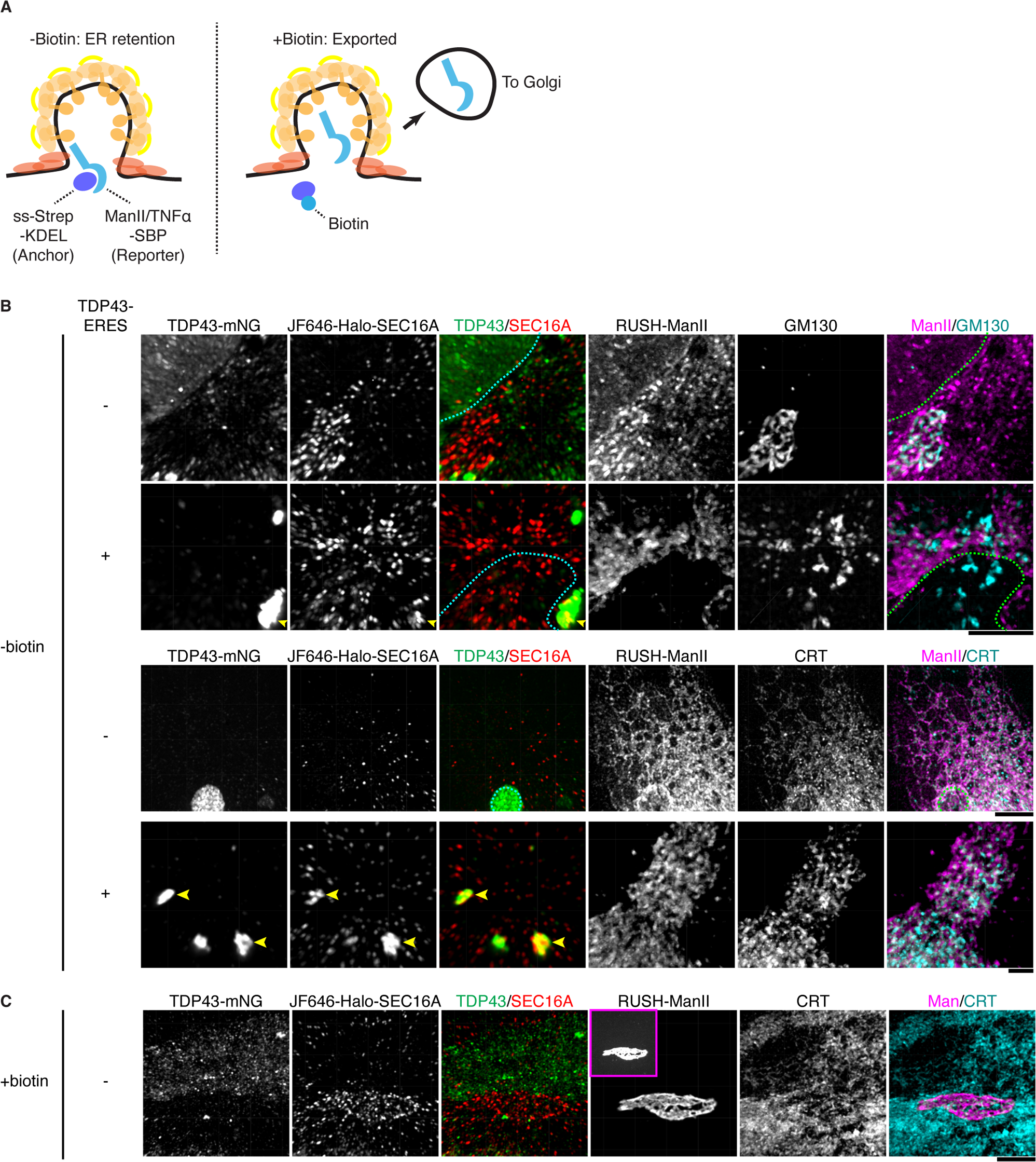
ER-to-Golgi transport of RUSH-ManII in cells without or with TDP43-ERES. (**A**) Working scheme of RUSH assay for examining ER-to-Golgi protein transport. Left: when cells are cultured in biotin-free media, the RUSH reporter containing streptavidin-binding protein (SBP) is retained in the ER or ERES through binding to the ER anchor ss-streptavidin-KDEL; Right: after biotin supplement, the reporter is released from anchor and subsequently exported to the Golgi. (**B**) Representative images of RUSH assay in cells without or with TDP-ERES. Cells cultured in biotin-free media was imaged with GM130 (Golgi) or Calreticulin (CRT – ER marker) co-stained by immunofluorescence. Arrowheads: TDP43-ERES; cyan or green dashes: the nuclei. The quantification of is shown in (Figure 5B and C). Each scale bar represents 5 μm. (**C**) Representative images of RUSH assay in cells without TDP-ERES and supplemented with biotin for 1 hr. CRT was co-stained to visualize the ER. The inset in the ManII channel shows a contrast-adjusted view.

**Figure S8.**
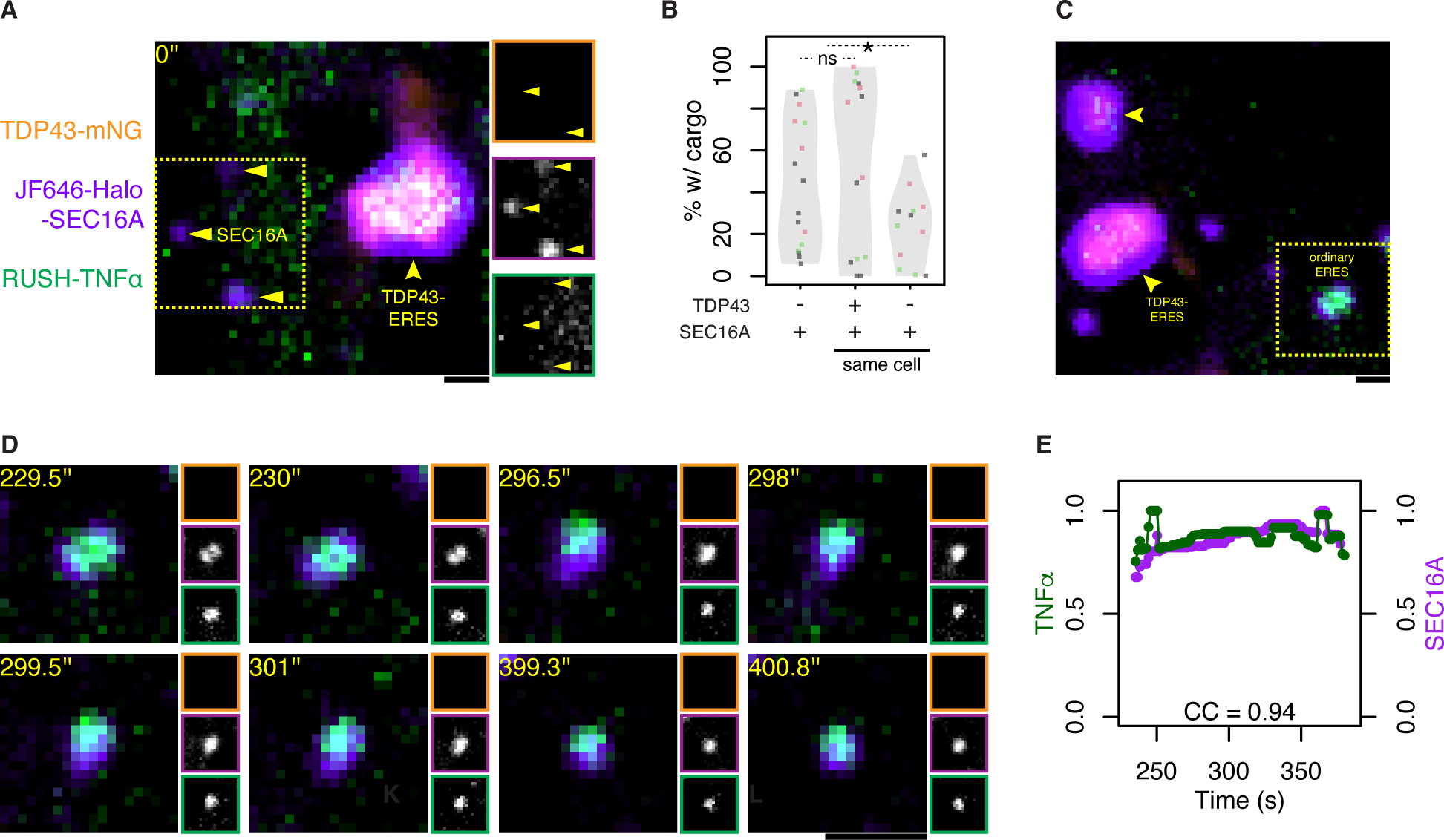
TDP43-ERES exhibit dominant effects over ordinary ERES in the same cell. (**A**) Zoom-out view of the TDP43-ERES (indicated by the arrowhead) in Figure 5G showing that SEC16A inclusions without TDP43 (indicated by triangles) in its vicinity are devoid of RUSH-TNFα upon biotin addition (0’’). Insets show individual channels of the yellow boxed region. Each scale bar represents 1 μm. (**B**) Quantification of RUSH experiments in Figure 5E and G showing the percentage of TDP43-ERES and SEC16A inclusions without TDP43 in either the same or different cells that recruited RUSH-TNFα. (**C**) Zoom-out view of an ordinary ERES (yellow boxed region) showing TDP43-ERES nearby (indicated by arrowheads). The timelapse recording of this ordinary ERES is shown in (D). (**D**) Selected frames of the timelapse recording of the ERES boxed in (C) during RUSH-TNFα assay. (**E**) Quantification of RUSH-TNFα and SEC16A intensities over time (normalized to their respective max intensities) in the TDP43-ERES tracked in (D), as in Figure 5F.

**Figure S9.**
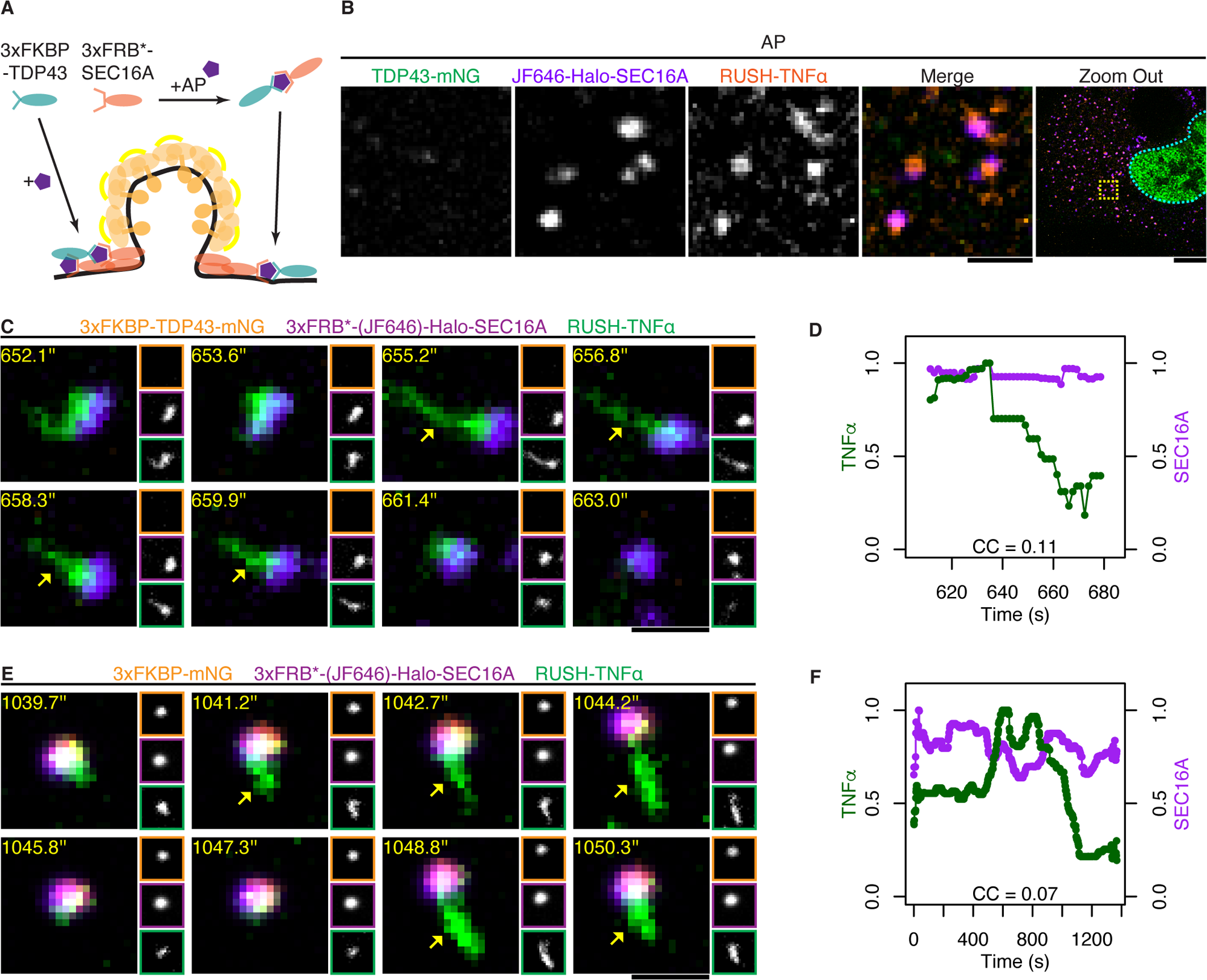
Controls experiments for AP21967-induced coaggregation of TDP43 with ERES. (**A**) Working scheme of AP21967-induced protein coaggregation. TDP43 (or mNG control) and SEC16A were respectively tagged with 3xFKBP and 3xFRB*, which can be induced to heterodimerize by both binding to the rapamycin analog AP21967 (AP). As such, AP induced aggregation of 3xFKBP-TDP43 at ERES marked by 3xFRB*-SEC16A. (**B**) Representative images of cells expressing TDP43-mNG, Halo-SEC16A and RUSH-TNFα and treated with AP21967 for 6 - 10 hr. In “Zoom Out”, the yellow box marks the region displayed in zoom-in views, and cyan dashes demarcate the nucleus. Scale bars except in “Zoom Out” represent 1 μm, and represent 5 μm in “Zoom Out”. (**C**) Selected frames of a representative timelapse recording of an ERES marked by 3xFRB*-(JF646)-Halo-SEC16A but not 3xFKBP-TDP43-mNG during RUSH-TNFα assay. The cell was not treated with AP21967. Arrows indicate the budding of a tubular transport intermediate. (**D**) Quantification of RUSH-TNFα and SEC16A intensities over time (normalized to their respective max intensities) in the TDP43-ERES tracked in (C). (**E**) Selected frames of a representative timelapse recording of an AP-induced mNG-ERES coagg during RUSH-TNFα assay. Arrows indicate the budding of tubular transport intermediates. (**F**) Quantification of RUSH-TNFα and SEC16A intensities over time (normalized to their respective max intensities) in the mNG-ERES tracked in (E).

**Figure S10.**
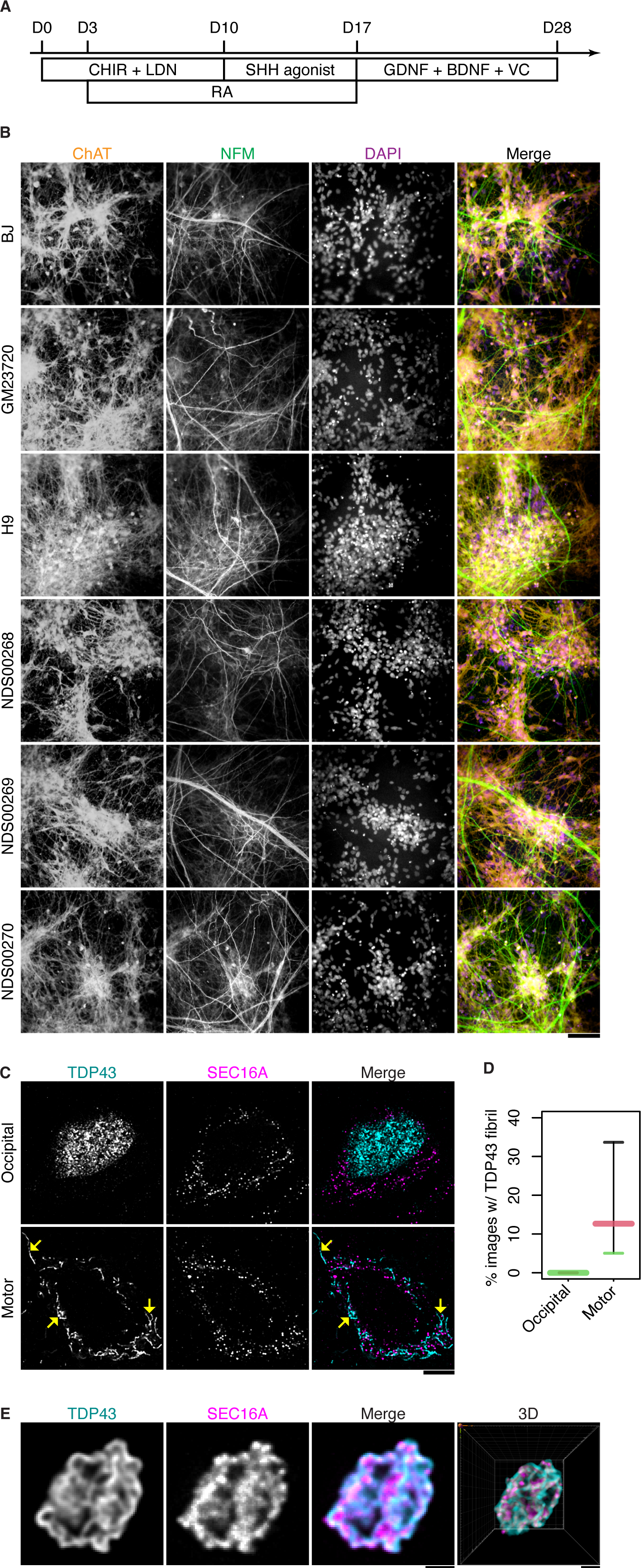
Neurons derived from stem cells or in postmortem brain sections of ALS patients. (**A**) Workflow of induced motor neuron (iMN) production from hESC/hiPSC. CHIR: CHIR99021; LDN: LDN193189; RA: retinoic acid; SHH: Sonic Hedgehog. VC: Vitamin C (L-ascorbic acid). D0: Day 0 of induced differentiation. (**B**) Representative images of the immunofluorescence staining of (cholinergic) neuronal markers ChAT and NFM in iMNs derived from non-ALS (BJ, GM23720 and H9) and ALS patient (NDS00268, NDS00269 and NDS00270) hESC/hiPSCs. Each scale bar represents 50 μm. (**C**) Representative images of the immunofluorescence staining of TDP43, SEC16A and NeuN in the post-mortem sections of the occipital cortex and motor cortex from the same ALS patient. Like Figure 7C and E but showing the presence of TDP43 fibrils (indicated by arrows) in the motor neurons. (**D**) Quantification of the experiments as in (C) comparing the percentage of images containing TDP43 fibrils in the occipital versus motor cortex of 3 different ALS patients (color-coded). At least 30 images were taken for each patient (*i.e.* N ≥ 3 x 30). (**E**) Zoom-in views of the curled TDP43/SEC16A skein in Figure 7E marked by the arrowhead.

**Video S1. TDP43 partitions dynamically between SGs and coaggregates with SEC16A.**

A representative movie of SGs and SEC16A inclusions in a cell shifted to 42°C for 60 min before recording started. Green: TDP43-mNG; Magenta: mCherry-G3BP1; Red: JF646-Halo-SEC16A. Annotated key frames and quantification are shown in Figure S4D and E, respectively.

**Video S2. TDP43 aggregates at ERES.**

A representative movie of ERES in a cell shifted to 42°C for 60 min before recording started. Cyan: TDP43-mNG; Magenta: JF646-Halo-SEC16A. Annotated key frames and quantification are shown in Figure 3C and D, respectively.

**Video S3. TDP43-ERES remain ER-associated.**

A representative movie of the movement of TDP43-ERES and SEC16A inclusions without TDP43 relative to the ER network. Green: TDP43-mNG; Purple: JF646-Halo-SEC16A; Orange: ss-mCherry-KDEL (ER). An annotated frame and quantification are shown in Figure 3I and J, respectively.

**Video S4. ERES prefer nascent over mature TDP43.**

A representative movie showing ERES and nucleus in a cell that was pre-stressed at 42°C for 6 hr and supplemented with 1 μM MG132 just before imaging started. Pre-existing (mature) Halo-TDP43 was pulse-labelled by JF646 (orange) while newly synthesized (new) Halo-TDP43 was labelled continuously with JF549 (green), and mEGFP-SEC16A (purple) was used to mark ERES. The workflow is illustrated in Figure S5E. Annotated key frames and quantification are shown in Figure S5F and G, respectively.

**Video S5. Partial FRAP of TDP43-ERES.**

A representative movie of a TDP43-ERES coagg after photobleaching in part by laser. Cyan: TDP43-mCherry; Magenta: mEGFP-SEC16A. Annotated key frames and quantification are shown in Figure 4A and B, respectively.

**Video S6. Partial FRAP of SG.**

A representative movie of a TDP43-containing SG after photobleaching in part by laser. Cyan: TDP43-mNG; Magenta: mCherry-G3BP1. Annotated key frames and quantification are shown in Figure S6A and B, respectively.

**Video S7. Full FRAP of TDP43-ERES.**

A representative movie of a TDP43-ERES coagg after photobleaching in its entirety by laser. Cyan: TDP43-mCherry; Magenta: mEGFP-SEC16A. Annotated key frames and quantification are shown in Figure 4D and E, respectively.

**Video S8. Full FRAP of SG.**

A representative movie of a TDP43-containing SG after photobleaching in its entirety by laser. Cyan: TDP43-mNG; Magenta: mCherry-G3BP1. Annotated key frames and quantification are shown in Figure S6C and D, respectively.

**Video S9. 1,6-hexanediol induces TDP43/SEC16A coaggregates.**

A movie showing a cell forming TDP43/SEC16A coaggs during 1,6-hexanediol (Hex) treatment. Cyan: TDP43-mCherry; Magenta: mEGFP-SEC16A. Hex was added between 7 min and 9 min. The 3D projection (top view) of an z stack is shown.

**Video S10. ERES without TDP43 export RUSH-TNFα.**

A representative movie of ERES without TDP43 exporting RUSH-TNFα after biotin supplement (at 0 sec). Orange: TDP43-mNG; Purple: JF646-Halo-SEC16A; Green: RUSH-TNFα. SEC16A inclusions were identified by ilastik (magenta contours) and tracked using Trackmate. Only the trajectory of the ERES shown in Figure 5E and F is displayed with color coding for time (blue: track start; red: track end). A tubule-like transport intermediate formed and released around 526 sec.

**Video S11. TDP43-ERES trap RUSH-TNFα.**

A representative movie of TDP43-ERES not exporting RUSH-TNFα after biotin supplement (at 0 sec). Orange: TDP43-mNG; Purple: JF646-Halo-SEC16A; Green: RUSH-TNFα. SEC16A inclusions were identified by ilastik (magenta contours) and tracked using Trackmate. Only the trajectory of the ERES shown in Figure 5G and H is displayed with color coding for time (blue: track start; red: track end).

**Video S12. ERES in the vicinity of TDP43-ERES trap RUSH-TNFα.**

A representative movie of ERES without TDP43 but in the vicinity of TDP43-ERES not exporting RUSH-TNFα after biotin supplement (at 0 sec). Orange: TDP43-mNG; Purple: JF646-Halo-SEC16A; Green: RUSH-TNFα. SEC16A inclusions were identified by ilastik (magenta contours) and tracked using Trackmate. Only the trajectory of the ERES shown in Figure S8D and E is displayed with color coding for time (blue: track start; red: track end).

**Video S11. AP21967-induced TDP43-ERES trap RUSH-TNFα.**

A representative movie of AP21967-induced TDP43-ERES not exporting RUSH-TNFα after biotin supplement (at 0 sec). Orange: 3xFKBP-TDP43-mNG; Purple: 3x-FRB*-(JF646)-Halo-SEC16A; Green: RUSH-TNFα. 3x-FRB*-(JF646)-Halo-SEC16A inclusions were identified by ilastik (magenta contours) and tracked using Trackmate. Only the trajectory of the ERES shown in Figure 6E and F is displayed with color coding for time (blue: track start; red: track end).

**Table S1. MS data.**

List of proteins identified in stress-induced aggregates and their Gene Ontology (GO) enrichment. (**Sheet 1**) The abundance of proteins identified by mass-spectrometry in the immunoprecipitate (IP) of aggregate-enriched sucrose fraction versus IP control. (**Sheet 2**) GO analysis of proteins enriched in aggregate by more than 8.5-fold compared with IP control showing the Biological Processes (BP) in which these proteins belong. (**Sheet 3**) GO analysis of proteins enriched in aggregate. Similar to (Sheet 2) but shows the Cellular Components (CC) where these proteins reside.

**Table S2. Plasmids, siRNAs and antibodies.**

Plasmids, DsiRNAs and antibodies used this this study. (**Sheet 1**) Plasmids. Transfection amounts of different plasmids were optimized for 35 mm glass-bottom dishes and using Lipofectamine 3000. Plasmids generated in this study and their maps and sequences will be submitted to Addgene and also available on request. (**Sheet 2**) The sense and antisense sequences of DsiRNAs against G3BP1 and G3BP2 and negative control DsiRNA (IDT NC1) which has no activity in human cells. The prefix “r” before A/U/C/G indicates ribonucleotide residues. (**Sheet 3**) Antibodies used in this study. Ig: immunoglobin.

## References

Alberti, S., Saha, S., Woodruff, J. B., Franzmann, T. M., Wang, J. & Hyman, A. A. 2018. A User’s Guide for Phase Separation Assays with Purified Proteins. J Mol Biol, 430, 4806–4820.w

Arzt, M., Deschamps, J., Schmied, C., Pietzsch, T., Schmidt, D., Tomancak, P., Haase, R. & Jug, F. 2022. LABKIT: Labeling and Segmentation Toolkit for Big Image Data. Frontiers in Computer Science, 4.

Balendra, R. & Isaacs, A. M. 2018. C9orf72-mediated ALS and FTD: multiple pathways to disease. Nat Rev Neurol, 14, 544–558.

Bastos, R., Ribas De Pouplana, L., Enarson, M., Bodoor, K. & Burke, B. 1997. Nup84, a novel nucleoporin that is associated with CAN/Nup214 on the cytoplasmic face of the nuclear pore complex. J Cell Biol, 137, 989–1000.

Beck, M. & Hurt, E. 2017. The nuclear pore complex: understanding its function through structural insight. Nat Rev Mol Cell Biol, 18, 73–89.

Bence, N. F., Sampat, R. M. & Kopito, R. R. 2001. Impairment of the ubiquitin-proteasome system by protein aggregation. Science, 292, 1552–5.

Berg, S., Kutra, D., Kroeger, T., Straehle, C. N., Kausler, B. X., Haubold, C., Schiegg, M., Ales, J., Beier, T., Rudy, M., Eren, K., Cervantes, J. I., Xu, B., Beuttenmueller, F., Wolny, A., Zhang, C., Koethe, U., Hamprecht, F. A. & Kreshuk, A. 2019. ilastik: interactive machine learning for (bio)image analysis. Nat Methods, 16, 1226–1232.

Bernad, R., Engelsma, D., Sanderson, H., Pickersgill, H. & Fornerod, M. 2006. Nup214-Nup88 nucleoporin subcomplex is required for CRM1-mediated 60 S preribosomal nuclear export. J Biol Chem, 281, 19378–86.

Biel, T. G., Aryal, B., Gerber, M. H., Trevino, J. G., Mizuno, N. & Rao, V. A. 2020. Mitochondrial dysfunction generates aggregates that resist lysosomal degradation in human breast cancer cells. Cell Death Dis, 11, 460.

Bodnar, A. G., Ouellette, M., Frolkis, M., Holt, S. E., Chiu, C. P., Morin, G. B., Harley, C. B., Shay, J. W., Lichtsteiner, S. & Wright, W. E. 1998. Extension of life-span by introduction of telomerase into normal human cells. Science, 279, 349–52.

Bolognesi, B., Faure, A. J., Seuma, M., Schmiedel, J. M., Tartaglia, G. G. & Lehner, B. 2019. The mutational landscape of a prion-like domain. Nat Commun, 10, 4162.

Boncompain, G., Divoux, S., Gareil, N., De Forges, H., Lescure, A., Latreche, L., Mercanti, V., Jollivet, F., Raposo, G. & Perez, F. 2012. Synchronization of secretory protein traffic in populations of cells. Nat Methods, 9, 493–8.

Bösl, B., Grimminger, V. & Walter, S. 2005. Substrate binding to the molecular chaperone Hsp104 and its regulation by nucleotides. J Biol Chem, 280, 38170–6.

Boutet, E., Lieberherr, D., Tognolli, M., Schneider, M. & Bairoch, A. 2007. UniProtKB/Swiss-Prot. Methods Mol Biol, 406, 89–112.

Budnik, A. & Stephens, D. J. 2009. ER exit sites--localization and control of COPII vesicle formation. FEBS Lett, 583, 3796–803.

Chiti, F. & Dobson, C. M. 2006. Protein misfolding, functional amyloid, and human disease. Annu Rev Biochem, 75, 333–66.

Chou, C.-C., Zhang, Y., Umoh, M. E., Vaughan, S. W., Lorenzini, I., Liu, F., Sayegh, M., Donlin-Asp, P. G., Chen, Y. H., Duong, D. M., Seyfried, N. T., Powers, M. A., Kukar, T., Hales, C. M., Gearing, M., Cairns, N. J., Boylan, K. B., Dickson, D. W., Rademakers, R., Zhang, Y.-J., Petrucelli, L., Sattler, R., Zarnescu, D. C., Glass, J. D. & Rossoll, W. 2018. TDP-43 pathology disrupts nuclear pore complexes and nucleocytoplasmic transport in ALS/FTD. Nature Neuroscience, 21, 228–239.

Consortium, T. G. O. 2015. Gene Ontology Consortium: going forward. Nucleic Acids Research, 43, D1049–D1056.

Copin, R., Baum, A., Wloga, E., Pascal, K. E., Giordano, S., Fulton, B. O., Zhou, A., Negron, N., Lanza, K. & Chan, N. 2021. The monoclonal antibody combination REGEN-COV protects against SARS-CoV-2 mutational escape in preclinical and human studies. Cell, 184, 3949–3961. e11.

Desantis, M. E., Leung, E. H., Sweeny, E. A., Jackrel, M. E., Cushman-Nick, M., Neuhaus-Follini, A., Vashist, S., Sochor, M. A., Knight, M. N. & Shorter, J. 2012. Operational plasticity enables hsp104 to disaggregate diverse amyloid and nonamyloid clients. Cell, 151, 778–793.

Düster, R., Kaltheuner, I. H., Schmitz, M. & Geyer, M. 2021. 1,6-Hexanediol, commonly used to dissolve liquid-liquid phase separated condensates, directly impairs kinase and phosphatase activities. J Biol Chem, 296, 100260.

Ellis, R. J. 2001. Macromolecular crowding: an important but neglected aspect of the intracellular environment. Curr Opin Struct Biol, 11, 114–9.

Ellis, R. J. & Minton, A. P. 2006. Protein aggregation in crowded environments. Biol Chem, 387, 485–97.

Erjavec, N., Larsson, L., Grantham, J. & Nyström, T. 2007. Accelerated aging and failure to segregate damaged proteins in Sir2 mutants can be suppressed by overproducing the protein aggregation-remodeling factor Hsp104p. Genes Dev, 21, 2410–21.

Ershov, D., Phan, M. S., Pylvänäinen, J. W., Rigaud, S. U., Le Blanc, L., Charles-Orszag, A., Conway, J. R. W., Laine, R. F., Roy, N. H., Bonazzi, D., Duménil, G., Jacquemet, G. & Tinevez, J. Y. 2022. TrackMate 7: integrating state-of-the-art segmentation algorithms into tracking pipelines. Nat Methods, 19, 829–832.

Frottin, F., Schueder, F., Tiwary, S., Gupta, R., Korner, R., Schlichthaerle, T., Cox, J., Jungmann, R., Hartl, F. U. & Hipp, M. S. 2019. The nucleolus functions as a phase-separated protein quality control compartment. Science, 365, 342–347.

Gallo, R., Rai, A. K., Mcintyre, A. B. R., Meyer, K. & Pelkmans, L. 2023. DYRK3 enables secretory trafficking by maintaining the liquid-like state of ER exit sites. Dev Cell, 58, 1880–1897.e11.

Gamerdinger, M., Kaya, A. M., Wolfrum, U., Clement, A. M. & Behl, C. 2011. BAG3 mediates chaperone-based aggresome-targeting and selective autophagy of misfolded proteins. EMBO Rep, 12, 149–56.

Garcia, D. M., Campbell, E. A., Jakobson, C. M., Tsuchiya, M., Shaw, E. A., Dinardo, A. L., Kaeberlein, M. & Jarosz, D. F. 2021. A prion accelerates proliferation at the expense of lifespan. Elife, 10.

Gates, S. N., Yokom, A. L., Lin, J., Jackrel, M. E., Rizo, A. N., Kendsersky, N. M., Buell, C. E., Sweeny, E. A., Mack, K. L., Chuang, E., Torrente, M. P., Su, M., Shorter, J. & Southworth, D. R. 2017. Ratchet-like polypeptide translocation mechanism of the AAA+ disaggregase Hsp104. Science, 357, 273–279.

Geiger, T., Wehner, A., Schaab, C., Cox, J. & Mann, M. 2012. Comparative proteomic analysis of eleven common cell lines reveals ubiquitous but varying expression of most proteins. Mol Cell Proteomics, 11, M111.014050.

Gleixner, A. M., Verdone, B. M., Otte, C. G., Anderson, E. N., Ramesh, N., Shapiro, O. R., Gale, J. R., Mauna, J. C., Mann, J. R., Copley, K. E., Daley, E. L., Ortega, J. A., Cicardi, M. E., Kiskinis, E., Kofler, J., Pandey, U. B., Trotti, D. & Donnelly, C. J. 2022. NUP62 localizes to ALS/FTLD pathological assemblies and contributes to TDP-43 insolubility. Nature Communications, 13, 3380.

Graham, F. L., Smiley, J., Russell, W. C. & Nairn, R. 1977. Characteristics of a human cell line transformed by DNA from human adenovirus type 5. J Gen Virol, 36, 59–74.

Grimminger-Marquardt, V. & Lashuel, H. A. 2010. Structure and function of the molecular chaperone Hsp104 from yeast. Biopolymers, 93, 252–76.

Halloran, M., Ragagnin, A. M. G., Vidal, M., Parakh, S., Yang, S., Heng, B., Grima, N., Shahheydari, H., Soo, K. Y., Blair, I., Guillemin, G. J., Sundaramoorthy, V. & Atkin, J. D. 2020. Amyotrophic lateral sclerosis-linked UBQLN2 mutants inhibit endoplasmic reticulum to Golgi transport, leading to Golgi fragmentation and ER stress. Cell Mol Life Sci, 77, 3859–3873.

Harari, A., Zoltsman, G., Levin, T. & Rosenzweig, R. 2022. Hsp104 N-terminal domain interaction with substrates plays a regulatory role in protein disaggregation. Febs j, 289, 5359–5377.

Harrison, A. F. & Shorter, J. 2017. RNA-binding proteins with prion-like domains in health and disease. Biochem J, 474, 1417–1438.

Hipp, M. S., Kasturi, P. & Hartl, F. U. 2019. The proteostasis network and its decline in ageing. Nat Rev Mol Cell Biol, 20, 421–435.

Hofmann, S., Kedersha, N., Anderson, P. & Ivanov, P. 2021. Molecular mechanisms of stress granule assembly and disassembly. Biochimica et Biophysica Acta (BBA) - Molecular Cell Research, 1868, 118876.

Hor, J. H., Santosa, M. M., Lim, V. J. W., Ho, B. X., Taylor, A., Khong, Z. J., Ravits, J., Fan, Y., Liou, Y. C., Soh, B. S. & Ng, S. Y. 2021. ALS motor neurons exhibit hallmark metabolic defects that are rescued by SIRT3 activation. Cell Death Differ, 28, 1379–1397.

Hor, J. H., Soh, E. S., Tan, L. Y., Lim, V. J. W., Santosa, M. M., Winanto, Ho, B. X., Fan, Y., Soh, B. S. & Ng, S. Y. 2018. Cell cycle inhibitors protect motor neurons in an organoid model of Spinal Muscular Atrophy. Cell Death Dis, 9, 1100.

Ilieva, E. V., Ayala, V., Jové, M., Dalfó, E., Cacabelos, D., Povedano, M., Bellmunt, M. J., Ferrer, I., Pamplona, R. & Portero-Otín, M. 2007. Oxidative and endoplasmic reticulum stress interplay in sporadic amyotrophic lateral sclerosis. Brain, 130, 3111–23.

Jackrel, M. E., Desantis, M. E., Martinez, B. A., Castellano, L. M., Stewart, R. M., Caldwell, K. A., Caldwell, G. A. & Shorter, J. 2014. Potentiated Hsp104 variants antagonize diverse proteotoxic misfolding events. Cell, 156, 170–82.

Jain, S., Wheeler, J. R., Walters, R. W., Agrawal, A., Barsic, A. & Parker, R. 2016. ATPase-Modulated Stress Granules Contain a Diverse Proteome and Substructure. Cell, 164, 487–98.

Jan, C. H., Williams, C. C. & Weissman, J. S. 2014. Principles of ER cotranslational translocation revealed by proximity-specific ribosome profiling. Science, 346.

Johnston, J. A., Ward, C. L. & Kopito, R. R. 1998. Aggresomes: a cellular response to misfolded proteins. J Cell Biol, 143, 1883–98.

Kawaguchi, Y., Kovacs, J. J., Mclaurin, A., Vance, J. M., Ito, A. & Yao, T. P. 2003. The deacetylase HDAC6 regulates aggresome formation and cell viability in response to misfolded protein stress. Cell, 115, 727–38.

Kayatekin, C., Matlack, K. E., Hesse, W. R., Guan, Y., Chakrabortee, S., Russ, J., Wanker, E. E., Shah, J. V. & Lindquist, S. 2014. Prion-like proteins sequester and suppress the toxicity of huntingtin exon 1. Proc Natl Acad Sci U S A, 111, 12085–90.

Khong, A., Matheny, T., Jain, S., Mitchell, S. F., Wheeler, J. R. & Parker, R. 2017. The Stress Granule Transcriptome Reveals Principles of mRNA Accumulation in Stress Granules. Mol Cell, 68, 808–820.e5.

Kon, T., Mori, F., Tanji, K., Miki, Y., Nishijima, H., Nakamura, T., Kinoshita, I., Suzuki, C., Kurotaki, H., Tomiyama, M. & Wakabayashi, K. 2022. Accumulation of Nonfibrillar TDP-43 in the Rough Endoplasmic Reticulum Is the Early-Stage Pathology in Amyotrophic Lateral Sclerosis. J Neuropathol Exp Neurol, 81, 271–281.

Kopito, R. R. 2000. Aggresomes, inclusion bodies and protein aggregation. Trends Cell Biol, 10, 524–30.

Kurokawa, K. & Nakano, A. 2019. The ER exit sites are specialized ER zones for the transport of cargo proteins from the ER to the Golgi apparatus. J Biochem, 165, 109–114.

Lim, L., Wei, Y., Lu, Y. & Song, J. 2016. ALS-Causing Mutations Significantly Perturb the Self-Assembly and Interaction with Nucleic Acid of the Intrinsically Disordered Prion-Like Domain of TDP-43. PLoS Biol, 14, e1002338.

Lin, Y., Protter, D. S., Rosen, M. K. & Parker, R. 2015. Formation and Maturation of Phase-Separated Liquid Droplets by RNA-Binding Proteins. Mol Cell, 60, 208–19.

Liu, B., Larsson, L., Caballero, A., Hao, X., Oling, D., Grantham, J. & Nystrom, T. 2010. The polarisome is required for segregation and retrograde transport of protein aggregates. Cell, 140, 257–67.

Mangiarotti, A., Chen, N., Zhao, Z., Lipowsky, R. & Dimova, R. 2022. Membrane wetting, molding and reticulation by protein condensates. bioRxiv, 2022.06.03.494704.

Mann, J. R., Gleixner, A. M., Mauna, J. C., Gomes, E., Dechellis-Marks, M. R., Needham, P. G., Copley, K. E., Hurtle, B., Portz, B., Pyles, N. J., Guo, L., Calder, C. B., Wills, Z. P., Pandey, U. B., Kofler, J. K., Brodsky, J. L., Thathiah, A., Shorter, J. & Donnelly, C. J. 2019. RNA Binding Antagonizes Neurotoxic Phase Transitions of TDP-43. Neuron, 102, 321–338.e8.

Marmor-Kollet, H., Siany, A., Kedersha, N., Knafo, N., Rivkin, N., Danino, Y. M., Moens, T. G., Olender, T., Sheban, D., Cohen, N., Dadosh, T., Addadi, Y., Ravid, R., Eitan, C., Toth Cohen, B., Hofmann, S., Riggs, C. L., Advani, V. M., Higginbottom, A., Cooper-Knock, J., Hanna, J. H., Merbl, Y., Van Den Bosch, L., Anderson, P., Ivanov, P., Geiger, T. & Hornstein, E. 2020. Spatiotemporal Proteomic Analysis of Stress Granule Disassembly Using APEX Reveals Regulation by SUMOylation and Links to ALS Pathogenesis. Mol Cell, 80, 876–891.e6.

Masrori, P. & Van Damme, P. 2020. Amyotrophic lateral sclerosis: a clinical review. Eur J Neurol, 27, 1918–1929.

Mateju, D., Franzmann, T. M., Patel, A., Kopach, A., Boczek, E. E., Maharana, S., Lee, H. O., Carra, S., Hyman, A. A. & Alberti, S. 2017. An aberrant phase transition of stress granules triggered by misfolded protein and prevented by chaperone function. Embo j, 36, 1669–1687.

Matsuki, H., Takahashi, M., Higuchi, M., Makokha, G. N., Oie, M. & Fujii, M. 2013. Both G3BP1 and G3BP2 contribute to stress granule formation. Genes Cells, 18, 135–46.

Matus, S., Valenzuela, V., Medinas, D. B. & Hetz, C. 2013. ER Dysfunction and Protein Folding Stress in ALS. Int J Cell Biol, 2013, 674751.

Medicherla, B. & Goldberg, A. L. 2008. Heat shock and oxygen radicals stimulate ubiquitin-dependent degradation mainly of newly synthesized proteins. J Cell Biol, 182, 663–73.

Medinas, D. B., Cabral-Miranda, F. & Hetz, C. 2019. ER stress links aging to sporadic ALS. Aging (Albany NY), 11, 5–6.

Medinas, D. B., Rozas, P., Martínez Traub, F., Woehlbier, U., Brown, R. H., Bosco, D. A. & Hetz, C. 2018. Endoplasmic reticulum stress leads to accumulation of wild-type SOD1 aggregates associated with sporadic amyotrophic lateral sclerosis. Proc Natl Acad Sci U S A, 115, 8209–8214.

Moda, F., Ciullini, A., Dellarole, I. L., Lombardo, A., Campanella, N., Bufano, G., Cazzaniga, F. A. & Giaccone, G. 2023. Secondary Protein Aggregates in Neurodegenerative Diseases: Almost the Rule Rather than the Exception. Front Biosci (Landmark Ed), 28, 255.

Neumann, M., Sampathu, D. M., Kwong, L. K., Truax, A. C., Micsenyi, M. C., Chou, T. T., Bruce, J., Schuck, T., Grossman, M., Clark, C. M., Mccluskey, L. F., Miller, B. L., Masliah, E., Mackenzie, I. R., Feldman, H., Feiden, W., Kretzschmar, H. A., Trojanowski, J. Q. & Lee, V. M. 2006. Ubiquitinated TDP-43 in frontotemporal lobar degeneration and amyotrophic lateral sclerosis. Science, 314, 130–3.

Nilsson, M. R. 2004. Techniques to study amyloid fibril formation in vitro. Methods, 34, 151–160.

Nishitoh, H., Kadowaki, H., Nagai, A., Maruyama, T., Yokota, T., Fukutomi, H., Noguchi, T., Matsuzawa, A., Takeda, K. & Ichijo, H. 2008. ALS-linked mutant SOD1 induces ER stress- and ASK1-dependent motor neuron death by targeting Derlin-1. Genes Dev, 22, 1451–64.

Nover, L., Scharf, K. D. & Neumann, D. 1989. Cytoplasmic heat shock granules are formed from precursor particles and are associated with a specific set of mRNAs. Mol Cell Biol, 9, 1298–308.

Ogrodnik, M., Salmonowicz, H., Brown, R., Turkowska, J., Sredniawa, W., Pattabiraman, S., Amen, T., Abraham, A. C., Eichler, N., Lyakhovetsky, R. & Kaganovich, D. 2014. Dynamic JUNQ inclusion bodies are asymmetrically inherited in mammalian cell lines through the asymmetric partitioning of vimentin. Proc Natl Acad Sci U S A, 111, 8049–54.

Oyanagi, K., Yamazaki, M., Takahashi, H., Watabe, K., Wada, M., Komori, T., Morita, T. & Mizutani, T. 2008. Spinal anterior horn cells in sporadic amyotrophic lateral sclerosis show ribosomal detachment from, and cisternal distention of the rough endoplasmic reticulum. Neuropathol Appl Neurobiol, 34, 650–8.

Park, S. H., Kukushkin, Y., Gupta, R., Chen, T., Konagai, A., Hipp, M. S., Hayer-Hartl, M. & Hartl, F. U. 2013. PolyQ proteins interfere with nuclear degradation of cytosolic proteins by sequestering the Sis1p chaperone. Cell, 154, 134–45.

Patel, A., Lee, H. O., Jawerth, L., Maharana, S., Jahnel, M., Hein, M. Y., Stoynov, S., Mahamid, J., Saha, S., Franzmann, T. M., Pozniakovski, A., Poser, I., Maghelli, N., Royer, L. A., Weigert, M., Myers, E. W., Grill, S., Drechsel, D., Hyman, A. A. & Alberti, S. 2015. A Liquid-to-Solid Phase Transition of the ALS Protein FUS Accelerated by Disease Mutation. Cell, 162, 1066–77.

Piovesan, D., Necci, M., Escobedo, N., Monzon, A. M., Hatos, A., Mičetić, I., Quaglia, F., Paladin, L., Ramasamy, P., Dosztányi, Z., Vranken, W. F., Davey, N. E., Parisi, G., Fuxreiter, M. & Tosatto, S. C. E. 2021. MobiDB: intrinsically disordered proteins in 2021. Nucleic Acids Res, 49, D361–d367.

Prasad, A., Bharathi, V., Sivalingam, V., Girdhar, A. & Patel, B. K. 2019. Molecular Mechanisms of TDP-43 Misfolding and Pathology in Amyotrophic Lateral Sclerosis. Front Mol Neurosci, 12, 25.

Reid, D. W. & Nicchitta, C. V. 2012. Primary role for endoplasmic reticulum-bound ribosomes in cellular translation identified by ribosome profiling. J Biol Chem, 287, 5518–27.

Robinson, J. L., Lee, E. B., Xie, S. X., Rennert, L., Suh, E., Bredenberg, C., Caswell, C., Van Deerlin, V. M., Yan, N., Yousef, A., Hurtig, H. I., Siderowf, A., Grossman, M., Mcmillan, C. T., Miller, B., Duda, J. E., Irwin, D. J., Wolk, D., Elman, L., Mccluskey, L., Chen-Plotkin, A., Weintraub, D., Arnold, S. E., Brettschneider, J., Lee, V. M. & Trojanowski, J. Q. 2018. Neurodegenerative disease concomitant proteinopathies are prevalent, age-related and APOE4-associated. Brain, 141, 2181–2193.

Rzechorzek, N. M., Thrippleton, M. J., Chappell, F. M., Mair, G., Ercole, A., Cabeleira, M., Rhodes, J., Marshall, I. & O’neill, J. S. 2022. A daily temperature rhythm in the human brain predicts survival after brain injury. Brain, 145, 2031–2048.

Saad, S. & Jarosz, D. F. 2021. Protein self-assembly: A new frontier in cell signaling. Curr Opin Cell Biol, 69, 62–69.

Saha, I., Yuste-Checa, P., Da Silva Padilha, M., Guo, Q., Körner, R., Holthusen, H., Trinkaus, V. A., Dudanova, I., Fernández-Busnadiego, R., Baumeister, W., Sanders, D. W., Gautam, S., Diamond, M. I., Hartl, F. U. & Hipp, M. S. 2023. The AAA+ chaperone VCP disaggregates Tau fibrils and generates aggregate seeds in a cellular system. Nat Commun, 14, 560.

Saito, K., Maeda, M. & Katada, T. 2017. Regulation of the Sar1 GTPase Cycle Is Necessary for Large Cargo Secretion from the Endoplasmic Reticulum. Front Cell Dev Biol, 5, 75.

Schindelin, J., Arganda-Carreras, I., Frise, E., Kaynig, V., Longair, M., Pietzsch, T., Preibisch, S., Rueden, C., Saalfeld, S., Schmid, B., Tinevez, J. Y., White, D. J., Hartenstein, V., Eliceiri, K., Tomancak, P. & Cardona, A. 2012. Fiji: an open-source platform for biological-image analysis. Nat Methods, 9, 676–82.

Scotter, E. L., Vance, C., Nishimura, A. L., Lee, Y. B., Chen, H. J., Urwin, H., Sardone, V., Mitchell, J. C., Rogelj, B., Rubinsztein, D. C. & Shaw, C. E. 2014. Differential roles of the ubiquitin proteasome system and autophagy in the clearance of soluble and aggregated TDP-43 species. J Cell Sci, 127, 1263–78.

Shi, Y., Lin, S., Staats, K. A., Li, Y., Chang, W. H., Hung, S. T., Hendricks, E., Linares, G. R., Wang, Y., Son, E. Y., Wen, X., Kisler, K., Wilkinson, B., Menendez, L., Sugawara, T., Woolwine, P., Huang, M., Cowan, M. J., Ge, B., Koutsodendris, N., Sandor, K. P., Komberg, J., Vangoor, V. R., Senthilkumar, K., Hennes, V., Seah, C., Nelson, A. R., Cheng, T. Y., Lee, S. J., August, P. R., Chen, J. A., Wisniewski, N., Hanson-Smith, V., Belgard, T. G., Zhang, A., Coba, M., Grunseich, C., Ward, M. E., Van Den Berg, L. H., Pasterkamp, R. J., Trotti, D., Zlokovic, B. V. & Ichida, J. K. 2018. Haploinsufficiency leads to neurodegeneration in C9ORF72 ALS/FTD human induced motor neurons. Nat Med, 24, 313–325.

Shorter, J. 2011. The mammalian disaggregase machinery: Hsp110 synergizes with Hsp70 and Hsp40 to catalyze protein disaggregation and reactivation in a cell-free system. PLoS One, 6, e26319.

Sontag, E. M., Samant, R. S. & Frydman, J. 2017. Mechanisms and Functions of Spatial Protein Quality Control. Annu Rev Biochem, 86, 97–122.

Soo, K. Y., Halloran, M., Sundaramoorthy, V., Parakh, S., Toth, R. P., Southam, K. A., Mclean, C. A., Lock, P., King, A., Farg, M. A. & Atkin, J. D. 2015. Rab1-dependent ER-Golgi transport dysfunction is a common pathogenic mechanism in SOD1, TDP-43 and FUS-associated ALS. Acta Neuropathol, 130, 679–97.

Soto, C. 2003. Unfolding the role of protein misfolding in neurodegenerative diseases. Nat Rev Neurosci, 4, 49–60.

Soto, C. & Pritzkow, S. 2018. Protein misfolding, aggregation, and conformational strains in neurodegenerative diseases. Nat Neurosci, 21, 1332–1340.

Spokoini, R., Moldavski, O., Nahmias, Y., England, J. L., Schuldiner, M. & Kaganovich, D. 2012. Confinement to organelle-associated inclusion structures mediates asymmetric inheritance of aggregated protein in budding yeast. Cell Rep, 2, 738–47.

Strambio-De-Castillia, C., Niepel, M. & Rout, M. P. 2010. The nuclear pore complex: bridging nuclear transport and gene regulation. Nat Rev Mol Cell Biol, 11, 490–501.

Suk, T. R. & Rousseaux, M. W. C. 2020. The role of TDP-43 mislocalization in amyotrophic lateral sclerosis. Mol Neurodegener, 15, 45.

Thul, P. J., Åkesson, L., Wiking, M., Mahdessian, D., Geladaki, A., Ait Blal, H., Alm, T., Asplund, A., Björk, L., Breckels, L. M., Bäckström, A., Danielsson, F., Fagerberg, L., Fall, J., Gatto, L., Gnann, C., Hober, S., Hjelmare, M., Johansson, F., Lee, S., Lindskog, C., Mulder, J., Mulvey, C. M., Nilsson, P., Oksvold, P., Rockberg, J., Schutten, R., Schwenk, J. M., Sivertsson, Å., Sjöstedt, E., Skogs, M., Stadler, C., Sullivan, D. P., Tegel, H., Winsnes, C., Zhang, C., Zwahlen, M., Mardinoglu, A., Pontén, F., Von Feilitzen, K., Lilley, K. S., Uhlén, M. & Lundberg, E. 2017. A subcellular map of the human proteome. Science, 356.

Tourrière, H., Chebli, K., Zekri, L., Courselaud, B., Blanchard, J. M., Bertrand, E. & Tazi, J. 2023. The RasGAP-associated endoribonuclease G3BP mediates stress granule assembly. J Cell Biol, 222.

Uversky, V. N. 2017. Intrinsically disordered proteins in overcrowded milieu: Membrane-less organelles, phase separation, and intrinsic disorder. Current Opinion in Structural Biology, 44, 18–30.

Vabulas, R. M. & Hartl, F. U. 2005. Protein synthesis upon acute nutrient restriction relies on proteasome function. Science, 310, 1960–3.

Vabulas, R. M., Raychaudhuri, S., Hayer-Hartl, M. & Hartl, F. U. 2010. Protein folding in the cytoplasm and the heat shock response. Cold Spring Harb Perspect Biol, 2, a004390.

Van Leeuwen, W., Nguyen, D. T. M., Grond, R., Veenendaal, T., Rabouille, C. & Farías, G. G. 2022. Stress-induced phase separation of ERES components into Sec bodies precedes ER exit inhibition in mammalian cells. J Cell Sci, 135.

Van Rheenen, W., Van Der Spek, R. A. A., Bakker, M. K., Van Vugt, J., Hop, P. J., Zwamborn, R. A. J., De Klein, N., Westra, H. J., Bakker, O. B., Deelen, P., Shireby, G., Hannon, E., Moisse, M., Baird, D., Restuadi, R., Dolzhenko, E., Dekker, A. M., Gawor, K., Westeneng, H. J., Tazelaar, G. H. P., Van Eijk, K. R., Kooyman, M., Byrne, R. P., Doherty, M., Heverin, M., Al Khleifat, A., Iacoangeli, A., Shatunov, A., Ticozzi, N., Cooper-Knock, J., Smith, B. N., Gromicho, M., Chandran, S., Pal, S., Morrison, K. E., Shaw, P. J., Hardy, J., Orrell, R. W., Sendtner, M., Meyer, T., Başak, N., Van Der Kooi, A. J., Ratti, A., Fogh, I., Gellera, C., Lauria, G., Corti, S., Cereda, C., Sproviero, D., D’alfonso, S., Sorarù, G., Siciliano, G., Filosto, M., Padovani, A., Chiò, A., Calvo, A., Moglia, C., Brunetti, M., Canosa, A., Grassano, M., Beghi, E., Pupillo, E., Logroscino, G., Nefussy, B., Osmanovic, A., Nordin, A., Lerner, Y., Zabari, M., Gotkine, M., Baloh, R. H., Bell, S., Vourc’h, P., Corcia, P., Couratier, P., Millecamps, S., Meininger, V., Salachas, F., Mora Pardina, J. S., Assialioui, A., Rojas-García, R., Dion, P. A., Ross, J. P., Ludolph, A. C., Weishaupt, J. H., Brenner, D., Freischmidt, A., Bensimon, G., Brice, A., Durr, A., Payan, C. A. M., Saker-Delye, S., Wood, N. W., Topp, S., Rademakers, R., Tittmann, L., Lieb, W., Franke, A., Ripke, S., Braun, A., Kraft, J., et al. 2021. Common and rare variant association analyses in amyotrophic lateral sclerosis identify 15 risk loci with distinct genetic architectures and neuron-specific biology. Nat Genet, 53, 1636–1648.

Wang, X., Zhou, X., Yan, Q., Liao, S., Tang, W., Xu, P., Gao, Y., Li, Q., Dou, Z., Yang, W., Huang, B., Li, J. & Zhang, Z. 2022. LLPSDB v2.0: an updated database of proteins undergoing liquid-liquid phase separation in vitro. Bioinformatics, 38, 2010–2014.

Watson, P., Townley, A. K., Koka, P., Palmer, K. J. & Stephens, D. J. 2006. Sec16 defines endoplasmic reticulum exit sites and is required for secretory cargo export in mammalian cells. Traffic, 7, 1678–87.

Weigel, A. V., Chang, C. L., Shtengel, G., Xu, C. S., Hoffman, D. P., Freeman, M., Iyer, N., Aaron, J., Khuon, S., Bogovic, J., Qiu, W., Hess, H. F. & Lippincott-Schwartz, J. 2021. ER-to-Golgi protein delivery through an interwoven, tubular network extending from ER. Cell, 184, 2412–2429.e16.

Weisberg, S. J., Lyakhovetsky, R., Werdiger, A. C., Gitler, A. D., Soen, Y. & Kaganovich, D. 2012. Compartmentalization of superoxide dismutase 1 (SOD1G93A) aggregates determines their toxicity. Proc Natl Acad Sci U S A, 109, 15811–6.

Wheeler, J. R., Matheny, T., Jain, S., Abrisch, R. & Parker, R. 2016. Distinct stages in stress granule assembly and disassembly. eLife, 5, e18413.

Yokom, A. L., Gates, S. N., Jackrel, M. E., Mack, K. L., Su, M., Shorter, J. & Southworth, D. R. 2016. Spiral architecture of the Hsp104 disaggregase reveals the basis for polypeptide translocation. Nat Struct Mol Biol, 23, 830–7.

Yu, G., Wang, L. G., Han, Y. & He, Q. Y. 2012. clusterProfiler: an R package for comparing biological themes among gene clusters. Omics, 16, 284–7.

Yuan, F., Alimohamadi, H., Bakka, B., Trementozzi, A. N., Day, K. J., Fawzi, N. L., Rangamani, P. & Stachowiak, J. C. 2021. Membrane bending by protein phase separation. Proc Natl Acad Sci U S A, 118.

Zacharogianni, M., Aguilera-Gomez, A., Veenendaal, T., Smout, J. & Rabouille, C. 2014. A stress assembly that confers cell viability by preserving ERES components during amino-acid starvation. Elife, 3.

Zappa, F., Wilson, C., Di Tullio, G., Santoro, M., Pucci, P., Monti, M., D’amico, D., Pisonero-Vaquero, S., De Cegli, R., Romano, A., Saleem, M. A., Polishchuk, E., Failli, M., Giaquinto, L. & De Matteis, M. A. 2019. The TRAPP complex mediates secretion arrest induced by stress granule assembly. Embo j, 38, e101704.

Zeskind, B. J., Jordan, C. D., Timp, W., Trapani, L., Waller, G., Horodincu, V., Ehrlich, D. J. & Matsudaira, P. 2007. Nucleic acid and protein mass mapping by live-cell deep-ultraviolet microscopy. Nat Methods, 4, 567–9.

Zhang, S., Cooper-Knock, J., Weimer, A. K., Shi, M., Moll, T., Marshall, J. N. G., Harvey, C., Nezhad, H. G., Franklin, J., Souza, C. D. S., Ning, K., Wang, C., Li, J., Dilliott, A. A., Farhan, S., Elhaik, E., Pasniceanu, I., Livesey, M. R., Eitan, C., Hornstein, E., Kenna, K. P., Blair, I., Wray, N. R., Kiernan, M., Mitne Neto, M., Chio, A., Cauchi, R., Robberecht, W., Van Damme, P., Corcia, P., Couratier, P., Hardiman, O., Mclaughin, R., Gotkine, M., Drory, V., Ticozzi, N., Silani, V., Veldink, J. H., Van Den Berg, L. H., De Carvalho, M., Mora Pardina, J. S., Povedano, M., Andersen, P., Weber, M., Başak, N. A., Al-Chalabi, A., Shaw, C., Shaw, P. J., Morrison, K. E., Landers, J. E., Glass, J. D., Veldink, J. H., Ferraiuolo, L., Shaw, P. J. & Snyder, M. P. 2022. Genome-wide identification of the genetic basis of amyotrophic lateral sclerosis. Neuron, 110, 992–1008.e11.

Zhou, C., Slaughter, B. D., Unruh, J. R., Eldakak, A., Rubinstein, B. & Li, R. 2011. Motility and segregation of Hsp104-associated protein aggregates in budding yeast. Cell, 147, 1186–96.

Zhou, C., Slaughter, Brian D., Unruh, Jay R., Guo, F., Yu, Z., Mickey, K., Narkar, A., Ross, Rhonda T., Mcclain, M. & Li, R. 2014. Organelle-Based Aggregation and Retention of Damaged Proteins in Asymmetrically Dividing Cells. Cell, 159, 530–542.

